# APOE 4/4 promotes dysfunctional and inflammatory phenotypes concomitant with impaired maturation of hiPSC-derived astrocytes

**DOI:** 10.64898/2026.09.04.749463

**Authors:** Rebeca Vecino, Eva Díaz-Guerra, Esther Arribas-González, David Sanz Gil, Francisco J. Fernández Acosta, Irene Serra, Elena P. Moreno-Jiménez, Rosario Moratalla, Marta Navarrete, Carlos Vicario

**Author notes:** **Corresponding author:** Carlos Vicario, Cajal Neuroscience Center, CSIC, Avenida de León 1, E-28805 Madrid, Spain. Rebeca Vecino and Eva Díaz-Guerra contributed equally to this article.

## Abstract

Alzheimer’s disease (AD) is the leading cause of dementia in the aging population, with the ε4 allele of apolipoprotein E (*APOE*) being the strongest genetic risk factor. Although astrocytes are a major source of APOE, how *APOE* alleles affect astrocyte maturation and function remains unclear. We generated human induced pluripotent stem cell (hiPSC)-derived astrocytes from AD patients carrying ε3/ε3 and ε4/ε4 alleles and from healthy controls (HC). We also used isogenic gene-edited hiPSC lines homozygous for each *APOE* allele and an *APOE* knock-out line to identify allele-specific phenotypes and distinguish gain-from loss-of-function mechanisms. APOE 4/4 astrocyte cultures showed significant reductions in GFAP- and S100β-positive cell percentages compared to APOE 2/2 and APOE 3/3, with no changes in GLT-1- and AQP4-positive cells. Astrocytes of all genotypes responded to IL-1β + TNFα by increasing proinflammatory cytokine expression and release, and to both IL-1β + TNFα and Aβ_1-42_ by changing morphology, with APOE 4/4 astrocytes showing increased *IL6* mRNA and morphological branching upon IL-1β + TNFα stimulation. Notably, under basal conditions, APOE 4/4 astrocytes showed significant reductions in glutamate uptake capacity and cell size alongside increased IL-6 release and *CXCL3* mRNA expression. In Aβ_1-42_ uptake experiments, the proportion of Aβ^+^astrocytes was higher in APOE 4/4 than in APOE KO cultures. Most phenotypes were absent in APOE KO astrocytes, suggesting that the effects of APOE 4/4 were predominantly mediated through gain-of-function mechanisms. Our results indicate that APOE 4/4 alters astrocyte morphological and molecular maturation while promoting inflammation, disturbing glutamate and Aβ handling under basal conditions. It suggests that APOE ε4/ε4 genotype disrupts astrocyte development and key processes of cellular homeostasis early in Alzheimer’s disease etiopathology.

## INTRODUCTION

Alzheimer’s disease (AD) is a progressive neurodegenerative disease characterized by the presence of extracellular deposits of beta-amyloid peptides (Aβ, forming the amyloid plaques), the intracellular accumulation of neurofibrillary tangles (NFTs) composed of hyperphosphorylated Tau and a chronic neuroinflammatory state (De Strooper & Karran 2016; Fernández-Calle et al., 2022; Ferrer, 2022; Frisoni et al., 2022; Gabitto et al., 2024; T. Guo et al., 2020; Hampel et al., 2021; Long & Holtzman, 2019; Rudman et al., 2026; Serrano-Pozo et al., 2021; Sierksma et al., 2020; van der Kant et al., 2020). Although age is a major risk factor for the development of neurodegenerative diseases, both genetic and environmental risk factors also play a determinant role in the onset and progression of AD. Autosomal dominant mutations in *APP* (Aβ precursor protein), *PSEN1* (presenilin1) and *PSEN2* (presenilin2), key players in the production and processing of amyloid peptides, cause fewer than 5% of all AD cases, known as early-onset AD (EOAD, also known as familial AD) [13,14]. Sporadic AD (late-onset AD or LOAD), the most common form of the disease, also exhibits an important genetic component, with the ε4 allele of *APOE* as the major risk factor [15–17]. Indeed, a recent study concluded that APOE ε4/ε4 homozygosity represents a genetic form of LOAD [18].

The human *APOE* gene encodes a ∼34 kDa glycoprotein, apolipoprotein E (APOE), a component of the main lipoprotein particles and the major cholesterol and lipid transporter in the central nervous system [19–21]. Among the different allelic variants (ε2, ε3 and ε4) that arise from two single-nucleotide polymorphisms in human *APOE* (rs7412 C/T and rs429358 C/T), the ε4 allele in homozygosis confers a 12- to 15-fold increase in disease risk, and may itself represent a genetic form of LOAD, as mentioned above [10,18,21–24]. Conversely, a protective role has been reported for the ε2 allele [25]. Although the *APOE* isoforms differ only in the amino acids at positions 112 and 158, these differences alter the APOE protein’s structure and modify its capacity to interact with Aβ, lipids and cholesterol, thereby impairing Aβ clearance and cholesterol transport [20,24,26].

The vast majority of brain APOE is produced by astrocytes, which are the primary source of APOE in the central nervous system; neurons and microglia also express *APOE* at lower basal levels, and neuronal *APOE* expression increases further with aging, stress, or injury [2,27,28]. Nonetheless, APOE protein and mRNA were reduced in APOE4 astrocytes, suggesting that the ε4 variant can negatively regulate its own transcription [29]. By contrast, a different study reported similar APOE levels in APOE3 and APOE4 hiPSC-derived astrocytes [30].

Beyond astrocytes, neuronal APOE4 has been reported to promote Tau phosphorylation and Aβ production [31] and to drive an early hippocampal excitation-inhibition imbalance during AD pathogenesis [31,32].

Astrocytes play a pivotal role in brain homeostasis, and are involved in critical functions such as neuronal differentiation and survival, modulation of synaptic transmission through glutamate uptake and release, maintenance of blood-brain barrier integrity, and regulation of bioenergetic and antioxidant defense (Abbott et al., 2006; Araque et al., 1999; Di Benedetto et al., 2022; Jiwaji & Hardingham, 2022; Linnerbauer et al., 2020; Verkhratsky & Nedergaard, 2018; Vicario-Abejón & Yusta-Boyo, 2004). Among these roles, astrocytic clearance of extracellular glutamate through the transporters GLT-1 (EAAT2) and GLAST (EAAT1) is essential to prevent excitotoxicity, and its impairment has been linked to synaptic dysfunction in AD [40]. Consequently, morphological, molecular and functional abnormalities in astrocytes contribute to the neurological decline observed in AD, in which *APOE* ε4/ε4 has been reported to affect several pathological processes, including Aβ seeding and accumulation together with subsequent reactive gliosis and neuroinflammation[7,17,37,40–43].

Reactive astrocytes typically undergo pronounced morphological remodeling — including changes in area, process length and branching complexity — that is increasingly used as a quantitative readout of astrocyte reactivity [40,44]. Importantly, these morphological changes are accompanied by functional alterations, particularly in the regulation and clearance of Aβ. Human iPSC-derived astrocytes carrying APOE 4/4 show impaired Aβ clearance and increased accumulation of Aβ oligomers compared with APOE 3/3 [29,45]. APOE4 astrocytes also exhibit a higher basal expression of inflammation-related genes, and an enhanced proinflammatory phenotype, and this APOE4 expression has been associated with chronic endoplasmic reticulum (ER) stress and mitochondrial dysfunction [45–50]. Consistent with a critical role of APOE ε4/ε4 in AD etiopathology, higher levels of Aβ oligomers have been reported in the brains of APOE ε4/ε4 carriers than in those of APOE ε3/ε3 carriers [18,51].

In recent years, an increasing number of studies employing stem cell technology to generate *in vitro / ex vivo* models of neurodegenerative diseases have demonstrated their strong potential to capture human processes and pathological conditions, including AD. Indeed, hiPSC-derived neurons, astrocytes, microglia-like cells and organoids recapitulate phenotypes associated with this disease [29,31,45,47,49,52–59].

Although these studies indicate progress in understanding the role of *APOE* polymorphism, and of *APOE* ε4/ε4 specifically in astrocytes in the context of AD, the cellular and molecular phenotypes and the mechanisms underlying cellular dysfunction remain to be fully elucidated. Therefore, further studies addressing *APOE*-related pathology (specifically that associated with *APOE* ε4/ε4) are needed to establish its correlation with AD etiopathology, progression and outcome, and to determine whether this relationship is causal.

Here, we developed a differentiation protocol to generate astrocytes from hiPSCs under defined culture-medium conditions in order to investigate the impact of AD and *APOE* polymorphism on astrocyte maturation, function, inflammatory response, morphology and Aβ uptake. Using both non-isogenic [60,61] and isogenic hiPSC lines [62], we found that APOE 4/4 alters astrocyte morphological and molecular maturation, reduces glutamate uptake, promotes IL-6 release and impairs Aβ handling under basal conditions. Furthermore, APOE 4/4 shapes the inflammatory and morphological profile of astrocytes in response to IL-1β + TNFα and to Aβ_1-42_. Our findings suggest that APOE 4/4 alters astrocyte maturation and function, with its effects emerging during astrocyte development in the context of AD.

## MATERIALS AND METHODS

### Growth and expansion of human iPSCs

Two different sets of vector-free hiPSC lines were used in this study. Four non-isogenic cell lines derived from patients with AD carrying APOE ε3 (two lines) and ε4 alleles (two lines) in homozygosis and two cell lines from HC were previously generated and characterized in our laboratory [60,61,63] and deposited in the Spanish National Stem Cell Bank/Banco Nacional de Líneas Celulares (BNLC, Instituto de Salud Carlos III, Spain). Once formed, the hiPSC colonies were mechanically divided into cell clumps, seeded onto mitotically inactivated mouse embryonic fibroblasts (MEFs), and expanded in hiPSC medium (KnockOut DMEM/F12, 100 μM NEAA, 2 mM GlutaMAX, 55 μM β-mercaptoethanol, 100 U/mL Penicillin, 100 μL/mL Streptomycin B, and 20% KnockOut serum replacement; ThermoFisher). Moreover, three isogenic hiPSC lines homozygous for the main *APOE* variants (ε2, ε3, and ε4) and an *APOE* knockout line (KO) were generated and characterized previously [62,64]. These hiPSCs were grown and expanded on vitronectin (VTN-N, Gibco™, ThermoFisher, A14700)-coated plates with Essential 8™ Medium (E8, Gibco™, ThermoFisher, A1517001). Cells were then washed with DPBS (Gibco™, ThermoFisher, 14190-094) and incubated with 0.5 mM EDTA for 5 min in a cell-culture incubator at 37 °C / 5% CO_2_ for passage and expansion.

### Human iPSC differentiation to astrocytes

Non-isogenic hiPSC colonies were grown on MEF feeders as above mentioned, and dissociated into a single-cell suspension by incubating them with a cell detachment solution (Accutase™, STEMCELL Technologies, 07920) for 10 min in a cell-culture incubator at 37 °C / 5% CO_2_. The cells were seeded onto 0.1% gelatin-coated dishes for 1.5 h to isolate them from MEFs (MEFs attach to the gelatin, while hiPSCs remain in the supernatant). Suspended hiPSCs were then transferred to low-attachment plates containing DMEM-F12 (Gibco™, ThermoFisher, 31330038) with N-2 supplement (Gibco™, ThermoFisher, 17502048) and B-27™ supplement minus vitamin A (B-27, Gibco™, ThermoFisher, 12587010) medium (referred to as DMEM-F12/N2/B27) supplemented with 250 ng/µL Noggin (Peprotech, 120-10C), 5 µM A83 (Miltenyi Biotec, 130-105-333) and 10 µM of ROCK inhibitor Y-27632 (Y27, TOCRIS, 1254) to initiate neural induction and generate embryoid bodies (EBs; day 0) (Figure 1A).

**Figure 1.**
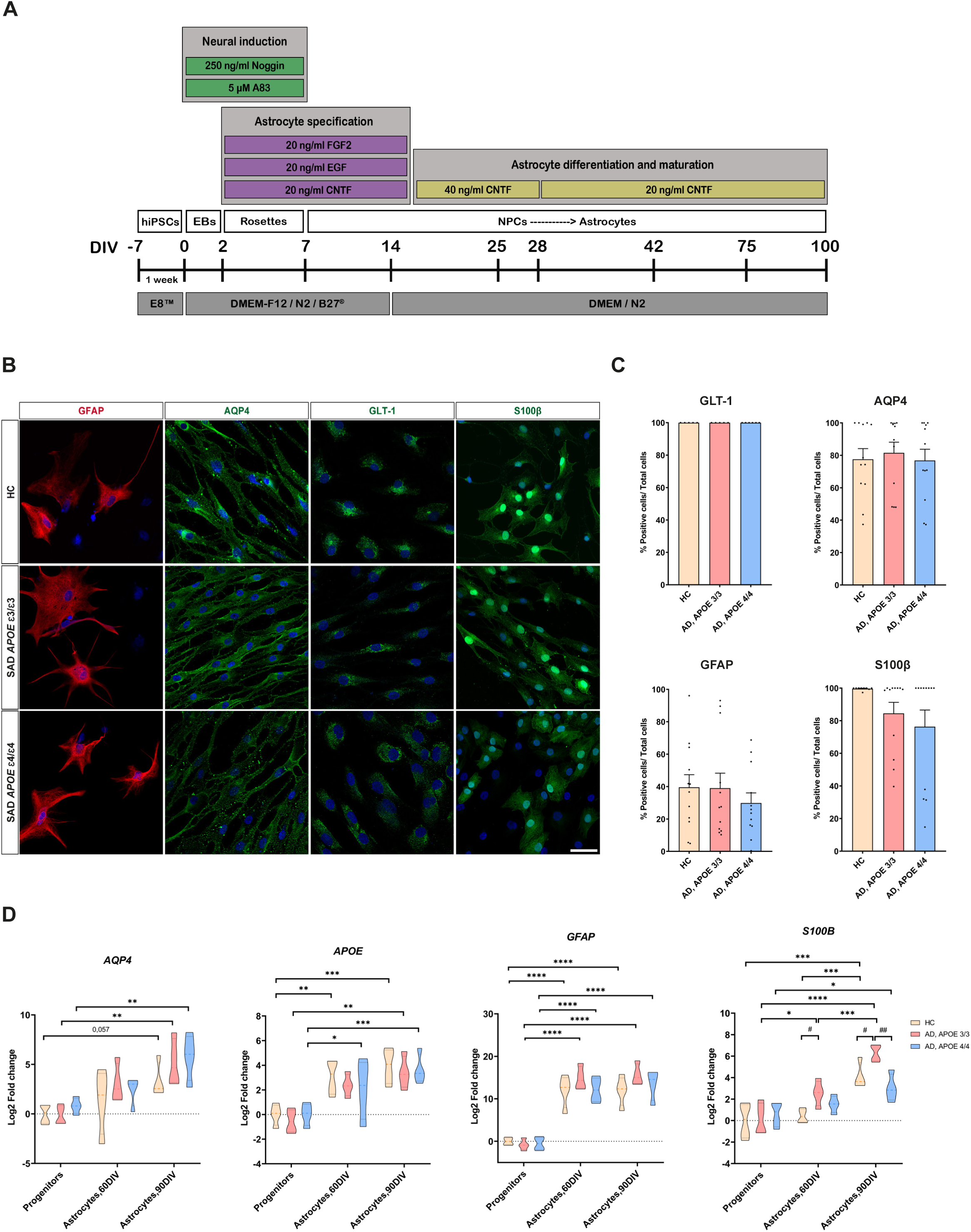
Human iPSC-derived astrocytes from HC and AD lines show no significant differences in astrocytic identity across *APOE* genotypes. **(A)** Schematic representation of the serum-free protocol for generating mature astrocytes from human iPSCs. **(B)** Representative immunocytochemistry images of astrocytic markers GFAP, AQP4, GLT-1, and S100β. Scale bar 50 μm. **(C)** Quantification of GLT-1^+^, AQP4^+^, GFAP^+^, and S100β^+^ astrocytes at 75-100 DIV. A non-significant reduction in the percentage of S100β^+^ cells was observed in AD, APOE 3/3 and AD, APOE 4/4 cultures compared with HC. One-way ANOVA with Tukey’s test or Kruskal-Wallis test followed by Dunn’s test. Results are mean ± SEM of n = 5-12 independent cultures/genotype. **(D)** RT-qPCR analysis of *AQP4*, *APOE, GFAP*, and *S100B* mRNA levels in hiPSC-derived neural progenitors (8-16 DIV) and astrocytes (60 and 90 DIV). While all markers were upregulated during differentiation-maturation, *S100B* exhibited a genotype-specific pattern at 90 DIV, with higher expression in AD, APOE 3/3 than in HC, but a marked decrease in AD, APOE 4/4 relative to AD, APOE 3/3. Statistical analysis was performed on ΔCt values using two-way ANOVA with Tukey’s test, and data were expressed as log_2_ fold change relative to the HC progenitor group. Asterisks (*) indicate differences between the time points within the same group. Hashtags (#) denote genotype differences at a given time point. \**p*<0.05, ^#^*p*<0.05, \*\**p*<0.01, ^##^*p*<0.01, \*\*\**p*<0.001, \*\*\*\**p*<0.0001. Results are mean ± SEM from n = 3 independent cultures/ genotype in technical triplicates. AD, Alzheimer’s disease; HC, healthy control.

*APOE* isogenic hiPSC colonies were grown on VTN-N, and 0.5 mM EDTA was applied for passage and expansion. To initiate differentiation, hiPSCs were incubated with accutase at 37 °C for 7 min, dissociated into single cells, and seeded into AggreWell800 plates (STEMCELL Technologies, 34811) at a density of 3 x 10^6^ single cells per well in E8 supplemented with 10 µM Y27. After 24 h, spheroids were collected from each microwell and transferred to low-attachment dishes containing DMEM-F12/N2/B27 medium supplemented with 250 ng/µL Noggin and 5 µM A83 to initiate neural induction and generate EBs (day 0).

After 2 days *in vitro* (DIV), isogenic and non-isogenic hiPSC-derived EBs were properly formed and passaged onto VTN-N-coated plates in DMEM-F12/N2/B27 medium supplemented with Noggin, A83, 20 ng/mL CNTF (Peprotech, 450-13), 20 ng/mL FGF2 (Peprotech, 100-18B), and 20 ng/mL EGF (Peprotech, AF-100- 15) (Figure 1A). On day 8, neural progenitor cells (NPCs) were incubated with accutase for 7 min and replated onto dishes coated with Cultrex^®^ ReadyBME (Bio-Techne, 3434-050-RTU) in DMEM-F12/N2/B27 medium supplemented with CNTF, EGF, and FGF2. On day 14, the medium was replaced with DMEM (Gibco™, ThermoFisher, 41966029) with N-2 supplement (referred to as DMEM/N2) supplemented with 40 ng/mL CNTF. After two weeks, the concentration of CNTF was reduced to 20 ng/mL. The medium was changed every 2-3 days until 90% cell confluence was reached, after which the cells were passaged and expanded or cryo-banked. On days 35-42, cells were seeded at a density of 20-25,000 cells/cm^2^ on Cultrex^®^- coated 12- and 24-well plates as well as on coated Thermanox™ coverslips in 24-well plates and maintained with DMEM/N2 medium supplemented with 20 ng/mL CNTF until they were used for experiments (see below). Non-isogenic hiPSCs were differentiated in parallel, and all assays and analyses were performed in parallel. The same strategy was used for the differentiation of isogenic hiPSCs and for the corresponding assays and analyses.

### Immunocytochemistry (ICC)

Astrocytes were grown for 75-90 DIV and fixed with 4% paraformaldehyde (PFA, PanReac AppliChem, 141451.1211) in PBS for 25 min at room temperature (RT). After washing three times with PBS, cells were permeabilized with permeabilization/blocking solution (10% goat/ donkey serum and 0.1-0.2% Triton X-100 in PBS) for 1 h at RT and subsequently incubated with primary antibodies in the blocking solution overnight at 4°C. The following antibodies were used: mouse anti-AQP4 (1:75, Santa Cruz Biotechnology, Cat# sc32739, RRID:AB_626695), mouse anti-Calnexin (1:40, Santa Cruz Biotechnology, Cat# sc23954, RRID:AB_626783), mouse anti-GFAP (1:1000, Millipore, Cat# MAB360, RRID:AB_11212597), rabbit anti-GFAP (1:1000, Agilent, Cat# Z0334, RRID:AB_10013382), mouse anti-GLT-1 (1:50, Santa Cruz Biotechnology, Cat# sc-365634, RRID:AB_10844832), rabbit anti-S100β (1:1500, Swant Cat# 37, RRID:AB_2315304) and rabbit anti-S100β (1:100, Proteintech Cat# 15146-1-AP, RRID:AB_2254244). The cells were then washed three times with PBS and incubated with the respective Alexa-conjugated secondary antibodies in blocking solution at RT for 1 h. Nuclei were stained with 1 µg/mL Hoechst 33342 (Sigma-Aldrich, 861405) for 5 min at RT. After three washes with PBS, the coverslips containing the cells were mounted with Mowiol 4-88 (Calbiochem, 475904) on glass microscope slides. No specific staining was observed in cells incubated solely with secondary antibodies.

### Microscopy and cell counting

Confocal images were acquired using a Leica TCS SP-5 and Stellaris 8 microscopes (Leica Microsystems srl, Milan, Italy) with the Leica Application Suite X software. Images were captured in the z-plane every 1-2 µm with a 40X oil-immersion objective at a resolution of 1024 x 1024. Ten random fields from each coverslip were analyzed to determine the percentage of astrocytes expressing the markers of interest.

Mosaic images were composed using a Leica Thunder imager 3D live cell fluorescence microscope (Leica Microsystems srl), which provided images of the entire coverslip area for cell counting and normalization of glutamate and IL-6 levels. Image processing and analysis were performed using the FIJI software (ImageJ, NIH, USA; Schindelin et al., 2012).

For morphological and Aβ uptake analyses, random images of 5–10 astrocytes displaying a clearly defined and isolated morphology were acquired from each coverslip. In total, 8–10 coverslips derived from 3–4 independent differentiations were analysed (see below).

### Amyloid beta uptake assay and microscopy analysis

Astrocytes were grown on coverslips for 75-85 DIV and incubated with fluorochrome-conjugated Aβ (1-42) (HiLyte™ Fluor 488-labeled, AnaSpec, AS- 60479-01) at a final concentration of 250 nM at 37 °C / 5% CO_2_. The vehicle medium, consisting of 1% NH4OH (AnaSpec, AS-61322), served as the negative control. After 24 h, the cells were washed twice with PBS to remove excess non-internalized Aβ and fixed with 4% PFA for 25 min for further ICC with specific antibodies against S100β and calnexin. Confocal microscope images of astrocytes that picked up Aβ were processed using FIJI software. An *ROI* of the whole astrocyte was defined, and the threshold for Aβ_1-42_ labelling was adjusted to determine the percentage of Aβ-tagged area and the percentage of Aβ-positive astrocytes. The percentage of calnexin-tagged area and the degree of Aβ and calnexin colocalization were also measured.

### Morphological analysis and three-dimensional reconstruction (3-D image)

Morphological analysis was performed using FIJI software on GFAP- or S100β- labeled astrocytes, which provided precise visualization of cell shape and processes. After capturing images in the z-plane every 1 µm and generating a maximum projection, the astrocyte profile was traced and saved as a region of interest (*ROI)*. The following ROI parameters were analyzed using the *measure* command: area, perimeter, and solidity. Sholl analysis was performed using the simple neurite tracer (*SNT*) plugin [66] by tracing astrocyte processes and placing concentric circles around the center of the cell nucleus with radial increments of 4 µm. The total length and the number of primary and total processes were measured as well.

### Treatment with proinflammatory stimuli: IL-1β + TNFα and Aβ_1-42_ (Inflammatory assay)

Astrocytes (75-80 DIV) were exposed to two different protocols to induce an inflammatory response. After washing with DPBS (Gibco™, ThermoFisher, 14190-094), the cells were treated with 50 ng/mL human TNFα (Peprotech, 300- 01A) and 10 ng/mL human IL-1β (Peprotech, 200-01B) or with 4 µM amyloid-β protein (1-42) (Bachem, BA-01-4014447). As control conditions, cells were treated with solutions in which the compounds were dissolved (0.1% BSA in water in the IL-1β + TNFα experiments or 5.8% DMSO in Tris Base in the Aβ_1-42_ experiments). After 48 h of incubation, the supernatants were collected and flash-frozen for ELISA, and the cells were fixed for ICC. Cell extracts were collected for further gene expression analyses.

### RT-qPCR

Total RNA extraction from cultured astrocyte progenitors and mature astrocytes was performed using the NucleoSpin® RNA kit (Macherey-Nagel, 740955.50) following the manufacturer’s instructions, and Nanodrop One (Thermo Scientific) was used to determine RNA concentration. The reverse transcription reaction was carried out with SuperScript™ III Reverse Transcriptase (Invitrogen, ThermoFisher, 18080044) on a thermal cycler (Techne, TC-312), and relative gene expression levels were quantified by RT-qPCR in triplicate with appropriate controls. A quantity of 12 μL of the Power SYBR™ Green PCR Master mix (Applied Biosystems, ThermoFisher, 4367659) containing the primers at a final concentration of 0,3 μM was then mixed with 8 μL of the cDNA sample and added to each well of a 96-well plate. qPCRs experiments were conducted on the 7500 Real-Time PCR system (Applied Biosystems, Software v2.3) using the following program: 2 min at 50 °C, 10 min at 95 °C, 40 cycles of 15 s at 95 °C, 1 min at 60 °C and a subsequent cycle of 15 s at 95 °C and 1 min at 60 °C ramping up to 95 °C (0.5 °C every 30 s) to measure the dissociation curve.

Primer pairs were designed using Lasergene software (DNASTAR) and selected according to the least probability of amplifying non-specific products and high ΔG values to avoid primer dimers and hairpin formations (as verified by the NCBI Primer-BLAST software) (Supplementary Table S1). All RT-qPCR cDNA products were sequenced to verify that they corresponded to the expected fragments of interest. The amplification dynamic range for each set of primers was assessed by performing serial cDNA dilutions. All primers had 95-105% efficiency.

The comparative C_T_ method (ΔΔC_T_) with the equation 2^-ΔΔC^_T_ [67,68] was used to analyze quantitative gene expression (fold change), and *GAPDH* was selected as the housekeeping gene.

The ΔC_T_ values for each condition were used for statistical analysis, and the results were represented using violin plots after conversion to a linear scale (log_2_ of the fold change).

### Quantification of IL-6 in the culture medium

Astrocytes were grown on coverslips for 75 DIV and treated with human proinflammatory cytokines IL-1β and TNFα for 48 h or with the vehicle, as described above. The supernatant was transferred to Eppendorf tubes, and the cells were fixed for ICC studies (in order to obtain the total number of astrocytes). For the quantitative detection of IL-6 in the culture medium, a commercial human Uncoated ELISA kit was used according to the manufacturer’s guidelines (Invitrogen, ThermoFisher, 88-7066-22). When isogenic cells were studied, the final IL-6 concentration was normalized to the number of GLT-1 positive cells in each well, as determined by analyzing fluorescence microscopy mosaic images (see above).

### Calcium (Ca²⁺) imaging

Calcium levels in hiPSC-derived astrocytes were monitored by fluorescence microscopy using the green-fluorescent Ca^+2^ indicator Fluo-4 (Molecular Probes™, Invitrogen™, F14201). Astrocytes were grown on coverslips for 90- 100 DIV and incubated with fluo-4-AM (2-10 μM) for 15-20 min at 37 °C. Images were recorded using a CCD camera (Hamamatsu ORCA-R2 C10600) attached to an Olympus BX51WI upright microscope. Fluo-4 was excited at 470 nm using a CoolLED pE-100 light source with exposure times of 200–500 ms, and images were acquired every 0.5–1 s. The CoolLED and CCD camera were controlled and synchronized by the IP Lab software (BD Biosciences, MD, USA), which was also used for quantitative epifluorescence measurements. Initial processing of Ca²⁺ recordings was performed using Image J Software (public domain software developed at the US NIH). Minor XY drift in image stacks was corrected post hoc using TurboReg plugin. For each recording, individual regions of interest (ROIs) were manually defined around single astrocytes, and mean fluorescence intensity within each ROI was calculated over time. Ca²⁺ signals were analyzed in MATLAB (R2018a; MathWorks, Natick, MA, USA) from raw fluorescence values as previously described [69]. Ca²⁺ responses were quantified under basal conditions and following mechanical stimulation. Mechanical stimulation was induced by gently touching an individual astrocyte with a glass micropipette. Three parameters were analyzed: mean Ca²⁺ event amplitude (ΔF/F₀), percentage of responding cells, and mean Ca²⁺ event frequency (events min⁻¹) for each genotype and experimental condition.

### Glutamate uptake assay

Astrocytes were grown in 24-well plates for 70-75 DIV and incubated with 40 μM glutamic acid (Sigma-Aldrich, Merk, G8415) diluted in Hank’s Balanced Salt Solution (HBSS; Gibco™, ThermoFisher, 14025092) for 30 min at 37 °C / 5% CO_2_. A culture medium containing 40 μM glutamic acid was used as the control for subsequent uptake analysis. The culture supernatant was collected, and the glutamate concentration in the medium was measured using a glutamate colorimetric assay kit (Sigma-Aldrich, ThermoFisher, MAK004), according to the manufacturer’s specifications. Glutamate uptake by astrocytes was determined by subtracting the amount of glutamate measured in the medium from the amount initially added to the assay. In all conditions, the final glutamate concentration was normalized to the number of GLT-1 positive cells in each well, determined by analyzing fluorescence microscopy mosaic images (see above), and expressed relative to HC or APOE 3/3 mean values.

### Statistical analysis

Data from astrocyte progenitors or from mature astrocytes collected at different DIV were combined for the analyses. All graphical representations and statistical analyses were conducted with GraphPad Prism 8 software (GraphPad, San Diego, CA, USA). The results are expressed as the mean ± standard error of the mean (SEM). All data points were plotted individually in each bar graph. Statistical significance was set at p < 0.05. Normality (Gaussian distribution) was assessed by the Shapiro-Wilk test, and equal variances were measured using Bartlett’s test. When an atypical deviation of a value was observed, the ROUT method (Q = 1-2%) was used to identify possible outliers. The unpaired t-test (parametric test) or the Mann-Whitney test (non-parametric test) was used to compare two groups. One-way ANOVA with post hoc Tukey’s multiple comparison test was performed when comparing more than two groups with a normal distribution. When the data did not follow a Gaussian distribution, the Kruskal-Wallis non-parametric test with post hoc Dunn’s multiple comparisons test was applied. A two-way ANOVA followed by Tukey’s test for multiple comparisons was performed to determine the effect of two variables on an outcome or dependent variable. The Mander’s overlap coefficient (MOC) was used to quantify the degree of colocalization between the fluorophores.

The number of biological replicates, p-values, and statistical methods used in each experiment are mentioned in the Results and/or figure legends.

## RESULTS

### *APOE* polymorphism differentially influences the maturation and marker expression of human iPSC-derived astrocytes

To investigate the impact of AD on human astrocyte development, we established a protocol to generate mature astrocytes from hiPSCs obtained from both AD patients and HC (Figure 1A) as detailed in Material and Methods. The experiments were performed using four non-isogenic cell lines from AD patients carrying *APOE ε3/ε3* and *ε4/ε4* alleles (designated as AD, APOE 3/3 or AD, APOE 4/4, respectively), along with two HC lines (*APOE ε3/ε3*), as mentioned above. Our protocol was implemented under serum-free conditions to avoid any impact of serum on cell biology and astrocyte phenotype [70]. We employed small molecules for dual SMAD inhibition to guide the hiPSCs towards a neuroectodermal fate, followed by the addition of growth and neurotrophic factors to promote cell proliferation and astrocyte differentiation and maturation. By adhering to this protocol, we achieved a nearly pure population of mature astrocytes at different *in vitro* differentiation stages.

To validate the efficiency of our differentiation protocol and confirm the astroglial identity of the generated cells, we characterized the non-isogenic astrocytes at different stages by cellular and molecular methods. First, to assess culture purity and maturity, immunocytochemistry (ICC) was performed at 75-90 DIV and canonical astrocyte markers such as GFAP, AQP4, GLT-1 and S100β were analyzed [71–74] (Figure 1B). Quantification of confocal images at this stage revealed a highly enriched population of astrocytes across all non-isogenic lines, with no statistically significant differences in the percentage of positive cells between HC, AD APOE 3/3 and AD APOE 4/4 lines (Figure 1C). Notably, the glutamate transporter GLT-1 and the water channel aquaporin-4 (AQP4) labeled close to 100% and 80% of cells, respectively, across all experimental groups, independent of the disease background. Although the percentage of GFAP- positive cells was below 50% in all cultures, the mature marker S100β labeled nearly all cells in HC condition, and showed a modest downward trend in AD APOE 3/3 (84.48%) and AD APOE 4/4 (76.35%) groups (Figure 1C).

We next asked whether the AD genetic background alters the temporal trajectory of astrogliogenesis, tracking the expression of canonical astrocyte transcripts and that of *APOE* across key developmental stages. *AQP4*, *APOE, GFAP*, and *S100B* were measured at the progenitor stage (8-16 DIV) and at 60 and 90 DIV by quantitative real-time PCR. Our data revealed a significant, time-dependent upregulation of these markers as differentiation progressed from progenitors to late mature stages (\**p*<0.05, ^#^*p*<0.05, \*\**p*<0.01, ^##^*p*<0.01, \*\*\**p*<0.001, \*\*\*\**p*<0.0001) (Figure 1D and Supplementary Figure S1A). This temporal increase followed a similar course in all lines, with no significant differences between HC and AD patients for *AQP4*, *APOE, and GFAP* transcripts (Figure 1D). *S100B* expression, however, followed a relatively different pattern. Although its mRNA increased during differentiation in every condition, expression levels were significantly higher in the AD APOE 3/3 line compared to HC at 60 DIV (^#^*p*<0.05) (Figure 1D). Moreover, at 90 DIV, this higher *S100B* expression in the AD APOE 3/3 was evident not only relative to the HC (^#^*p*<0.05) but also compared to the AD APOE 4/4 line (^##^*p*<0.01). These findings suggest that changes in *S100B* expression during human astrogliogenesis might depend on the specific *APOE* variants.

To evaluate this hypothesis and better investigate the influence of *APOE* polymorphism on astrocyte development, we derived mature astrocytes from *APOE* isogenic hiPSC lines (originating from a parental line with the *APOE ε4/ε4* genotype), which carried the *ε2/ε2*, *ε3/ε3* and *ε4/ε4* alleles, along with an *APOE* KO line. They were designated hereafter as APOE 2/2, 3/3, 4/4, and KO. These lines, which differ exclusively in their *APOE* allele while sharing the same genetic background [62,64], facilitate direct genotype comparisons in a controlled environment. Cells were differentiated into mature astrocytes following the same protocol previously described (Figure 1A) and subsequently analyzed by ICC (Figure 2A-B) and RT-qPCR (Figure 2C-D) for a similar panel of astrocytic markers. ICC analysis of isogenic *APOE* astrocytes revealed that GLT-1 immunoreactivity was present in nearly all cells across all genotypes, reaching values close to 100%, with no statistically significant differences between groups (Figures 2A-B). Similarly, AQP4-positive cells ranged from 78% to 92% across genotypes, with no significant differences observed. In contrast, GFAP immunoreactivity was significantly reduced in APOE 4/4 cultures compared to all other genotypes (APOE 2/2, APOE 3/3 and APOE KO; \**p*<0.05, \*\*\**p*<0.001). Consistent with this observation, S100β immunopositivity also showed a significant decrease in APOE 4/4 cultures relative to APOE 2/2 and APOE 3/3 (\*\**p*<0.01, \*\*\**p*<0.001). A decrease in S100β immunopositivity was also found in APOE KO cultures compared with APOE 3/3 (\**p*<0.05) (Figures 2A-B). To evaluate transcriptional changes associated with astrocyte maturation and *APOE* expression, RT-qPCR analysis was conducted at two timepoints: astrocyte progenitors (8-16 DIV) and astrocytes (60-75 DIV) (Figure 2C-D and Supplementary Figure S1B). Gene expression analysis revealed that *AQP4* and *S100B* mRNA levels increased significantly during astrocyte maturation across genotypes, with no differences between them at either timepoint for *AQP4*, confirming a largely consistent maturation trajectory regardless of *APOE* allele (\**p*<0.05, \*\**p*<0.01, \*\*\**p*<0.001, \*\*\*\**p*<0.0001) (Figure 2C and Supplementary Figure S1B). Furthermore, *S100B* expression also increased during astrocyte maturation across all genotypes, although it was significantly higher in APOE KO astrocytes compared to APOE 4/4 and APOE 2/2 (Figure 2C: ^#^*p*<0.05, ^##^*p*<0.01). For *APOE* and *SLC1A2*/*GLT1*, the increase during maturation reached statistical significance only in APOE 3/3 and APOE 4/4 astrocytes, respectively (\*\**p*<0.01, \*\*\*\**p*<0.0001), with no differences detected between genotypes at either stage (Figure 2C and Supplementary Figure S1B). Although *GFAP* mRNA expression could not be properly assessed in progenitor cells, no significant differences between genotypes were detected in mature astrocytes (Figure 2D).

**Figure 2.**
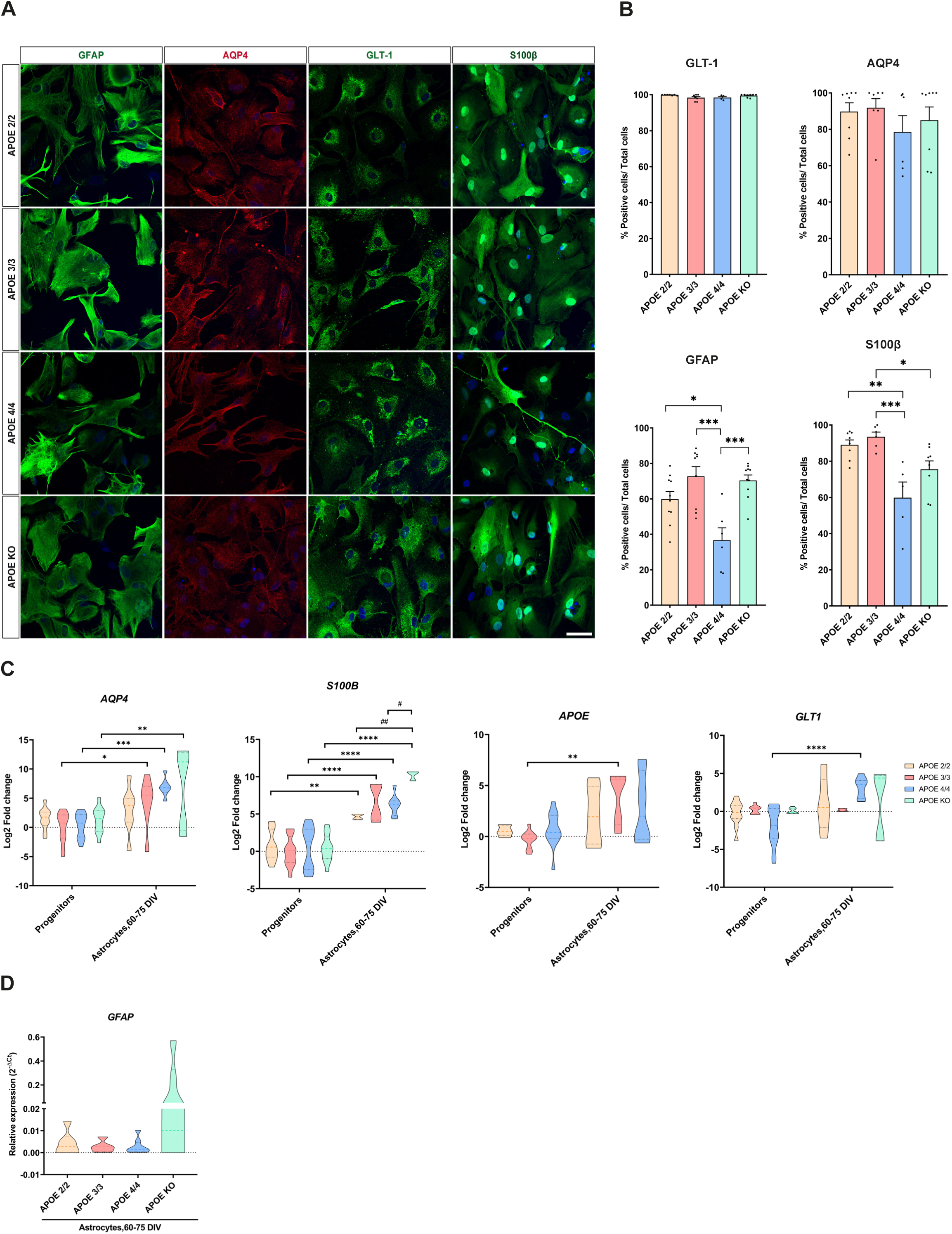
*APOE4* homozygosity reduces GFAP and S100β marker expression in isogenic human iPSC-derived astrocytes. **(A)** Representative immunocytochemistry images for GFAP, AQP4, GLT-1, and S100β in isogenic *APOE* cultures at 75-100 DIV. Scale bar 50 μm. **(B)** Quantification of GLT-1, AQP4, GFAP, and S100β markers. GFAP+ cells were significantly reduced specifically in APOE 4/4 cultures. S100β+ cells were significantly lower in APOE 4/4 than in APOE 2/2 and APOE 3/3 cultures, and in APOE KO than in APOE 3/3 cultures. Statistical significance was determined by one-way ANOVA with Tukey’s test, or Kruskal-Wallis test with Dunn’s test. \**p*<0.05, \*\**p*<0.01, \*\*\**p*<0.001. Results are mean ± SEM of n = 6-10 independent cultures per genotype. **(C)** mRNA expression of *AQP4*, *S100B*, *APOE,* and *SLC1A2*/*GLT1* in iPSC-derived neural progenitors (8-16 DIV) and astrocytes (60-75 DIV). All markers were upregulated upon differentiation, with the strongest increases in *AQP4* and *S100B* across genotypes, whereas *SLC1A2*/*GLT1* and *APOE* reached significance only in APOE 4/4 and APOE 3/3, respectively. At 60-75 DIV, *S100B* expression was significantly higher in APOE KO than in APOE 2/2 and APOE 4/4 astrocytes. Statistical significance was assessed on ΔCt values by two-way ANOVA with Tukey’s test; data are expressed as log_2_ fold change relative to APOE 3/3 progenitors. Asterisks (*) indicate differences between developmental stages within the same genotype; hashtags (#) denote differences between genotypes at a given stage. \**p*<0.05, ^#^*p*<0.05, \*\**p*<0.01, ^##^*p*<0.01, \*\*\**p*<0.001, \*\*\*\**p*<0.0001. Results are mean ± SEM from n = 4 independent cultures per genotype in technical triplicates. **(D)** Relative *GFAP* expression (2^⁻ΔCt^) in human astrocytes at 60–75 DIV across isogenic lines, showing no significant differences between genotypes. Statistical analysis was performed on ΔCt values using one- way ANOVA with Tukey’s test. Results are mean ± SEM from n = 4 independent cultures per genotype in technical triplicates.

Overall, these results indicate that APOE 4/4 cultures exhibit a distinct phenotypic profile marked by a selective reduction in GFAP- and S100β protein-expressing cells, while GLT-1 and AQP4 markers remain unaffected across genotypes. The discrepancy between mRNA and protein expression observed for both *GFAP* and *S100B* in APOE 4/4 astrocytes suggests that post-transcriptional mechanisms may contribute to their selective reduction at the protein level.

### *APOE4* homozygosity impairs glutamate uptake and promotes a basal proinflammatory astrocytic state

To evaluate the functional capabilities of astrocyte cultures and examine the impact of the *APOE* genotype on key astrocytic functions, both patient-derived and isogenic *APOE* astrocytes underwent a series of functional assays, such as glutamate uptake, Ca²⁺ signaling and inflammatory response (Figure 3, Figure 4, Supplementary Figure S2). Glutamate uptake was assessed at 70-75 DIV by ELISA, with values normalized to the HC and APOE 3/3 groups in patient-derived and isogenic cultures, respectively (Figure 3A). No clear differences were found between AD and HC groups, nor between AD APOE 3/3 and AD APOE 4/4 astrocytes. Nonetheless, there was a non-statistically significant trend toward reduced glutamate uptake in AD patients, particularly evident in AD APOE 4/4 cells. This observation was further supported by our isogenic model, in which APOE 4/4 astrocytes exhibited a statistically significant decrease in glutamate uptake compared to APOE 3/3 cells (***p<0.001), suggesting that *APOE4* homozygosity per se is sufficient to impair astrocytic glutamate clearance capacity.

**Figure 3.**
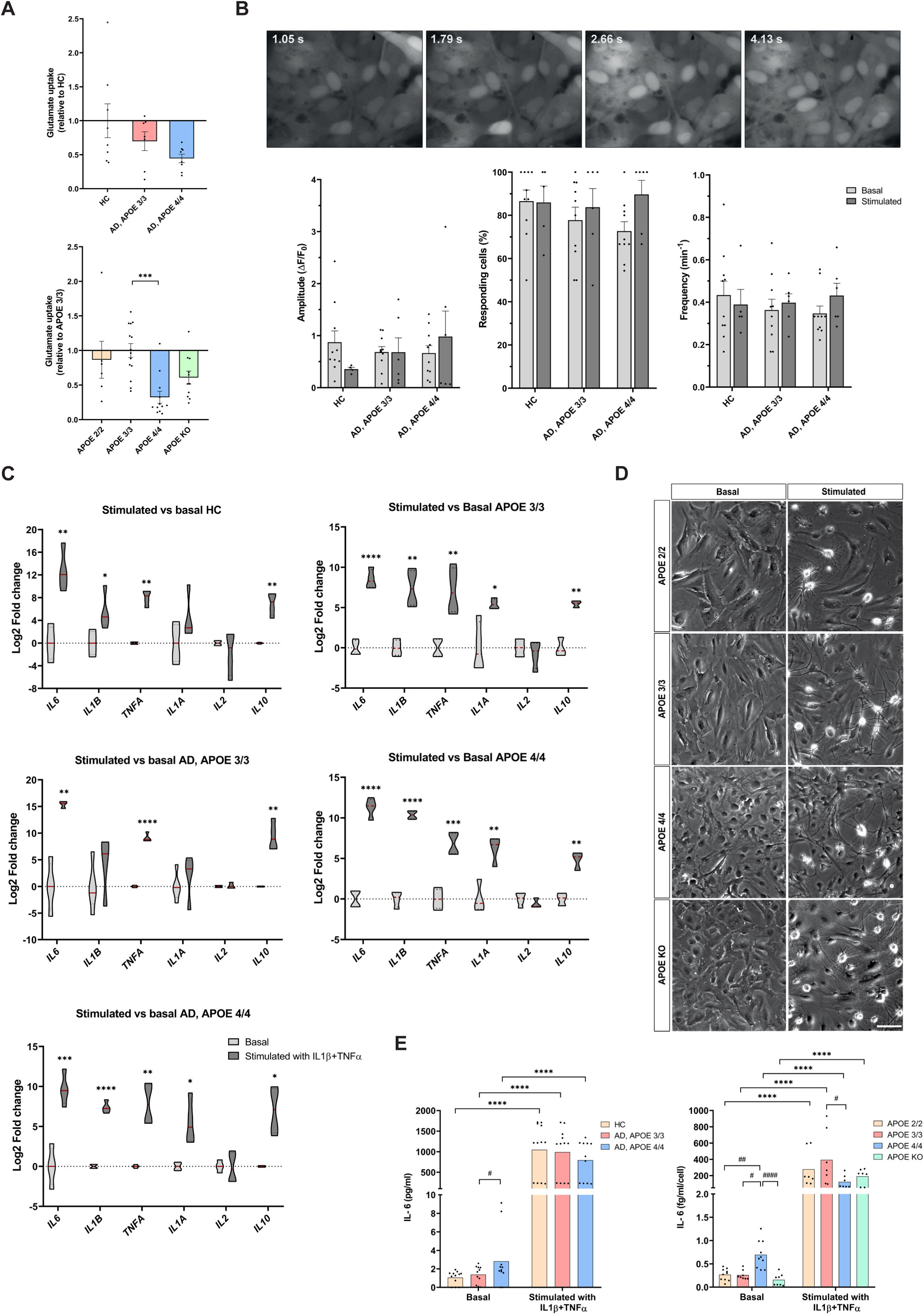
Glutamate uptake and IL-6 secretion are selectively dysregulated in APOE 4/4 astrocytes. **(A)** Functional glutamate clearance in HC and AD patient-derived (top) and isogenic *APOE* (bottom) astrocytes at 70-75 DIV. A downward trend in AD lines reached significance only in isogenic APOE 4/4 astrocytes. Data were normalized to HC or APOE 3/3, respectively. One-way ANOVA with Tukey’s test. \*\*\**p*<0.001. Results are mean ± SEM of n = 6-14 independent cultures/genotype. **(B)** Representative time-lapse images showing calcium wave propagation in HC- and AD-derived astrocytes at 90 DIV. Amplitude, responding-cell percentage, and frequency were determined using Fluo-4-AM. No significant differences were observed between genotypes or basal and stimulated conditions. Two-way ANOVA with Tukey’s test. Results are mean ± SEM of n = 6-10 independent cultures/genotype. **(C)** mRNA levels of proinflammatory (*IL6, IL1A, IL1B, TNFA, IL2*) and anti-inflammatory (*IL10*) cytokines in HC- and AD-derived (left) and isogenic APOE 3/3 and APOE 4/4 (right) astrocytes under basal and stimulated (IL-1β + TNFα) conditions. Most genes were upregulated upon stimulation, although responses varied; AD APOE 3/3 astrocytes showed a more restricted response (*IL6, TNFA, IL10*), while *IL2* was unchanged. Unpaired t-test on ΔCt values. \**p*<0.05, \*\**p*<0.01, \*\*\**p*<0.001, \*\*\*\**p*<0.0001. Results are mean ± SEM of n = 4 independent cultures/ genotype. **(D)** Representative phase-contrast images showing morphological remodeling after IL-1β + TNFα stimulation. Scale bar 100 μm. **(E)** Extracellular IL-6 determined by ELISA in patient-derived (left) and isogenic *APOE* (right) astrocytes at 75 DIV. Basal IL-6 secretion was higher in AD APOE 4/4 than APOE 3/3 and in isogenic APOE 4/4 than all other genotypes. Stimulation induced robust IL-6 secretion across groups, but levels were lower in isogenic APOE 4/4 than APOE 3/3. Two-way ANOVA with Tukey’s test. Asterisks (*) indicate basal versus stimulated differences; hashtags (#) denote genotype differences within condition. \**p*<0.05, ^#^*p*<0.05, \*\**p*<0.01, ^##^*p*<0.01, \*\*\**p*<0.001, ^###^*p*<0.001, \*\*\*\**p*<0.0001, ^####^*p*<0.0001. Results are mean ± SEM of n = 7-12 independent cultures/ genotype. AD, Alzheimer’s disease; HC, healthy controls.

**Figure 4.**
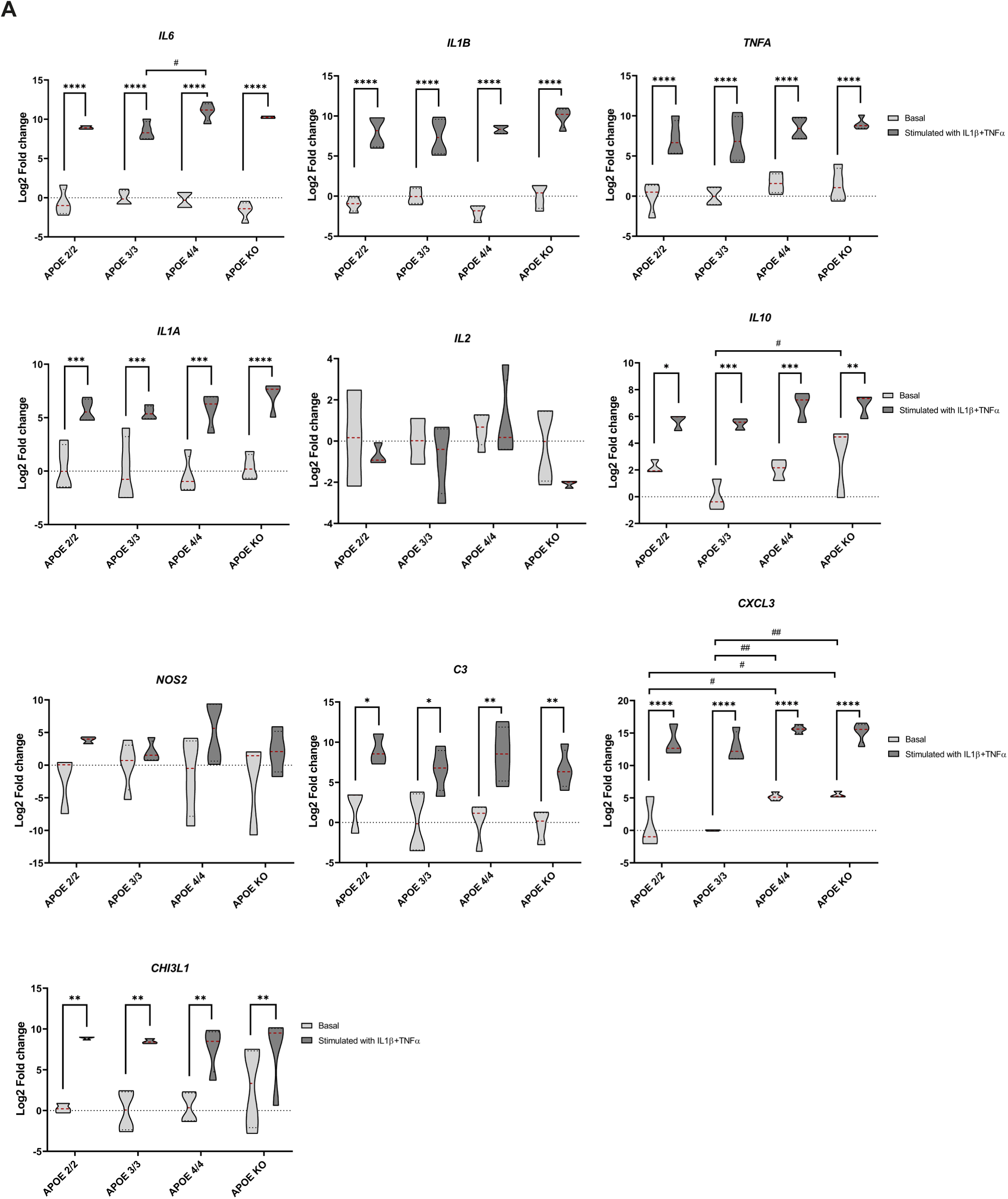
Genotype-specific inflammatory signatures emerge in *APOE*- isogenic astrocytes. mRNA expression of different cytokines and inflammatory markers was analyzed in human isogenic astrocytes (APOE ε2/ε2, ε3/ε3, ε4/ε4 and KO) at 75-90 DIV. Treatment with IL-1β + TNFα induced a significant increase in the mRNA expression of nearly all analyzed inflammatory markers across all genotypes, with the exception of *IL2* and *NOS2*, which were not significantly modulated in any genotype. Under basal conditions, APOE KO astrocytes showed significantly higher *IL10* expression than APOE 3/3 cells, whereas both APOE 4/4 and KO lines exhibited increased basal *CXCL3* levels relative to APOE 2/2 and APOE 3/3 astrocytes. Following stimulation, APOE 4/4 cells displayed significantly higher *IL6* upregulation than APOE 3/3. Two-way ANOVA was performed on ΔCt values, followed by Tukey’s test. Data are presented as log_2_ fold change relative to the APOE 3/3 basal condition. Asterisks (*) indicate differences between basal and stimulated conditions within genotype; hashtags (#) denote genotype differences within condition. \**p*<0.05, ^#^*p*<0.05, \*\**p*<0.01, ^##^*p*<0.01, \*\*\**p*<0.001, \*\*\*\**p*<0.0001. Results are mean ± SEM of n = 4 independent cultures/ genotype.

In addition, Ca²⁺ signaling dynamics were assessed in patient-derived astrocytes by live imaging under both spontaneous (basal) and mechanically evoked (stimulated) conditions. Sequential frames illustrating Ca²⁺ responses across the astrocyte network are shown in Figure 3B, with the complete recording available in the supplementary information (Supplementary Video S1). Ca²⁺ response parameters, including event amplitude, the proportion of responding cells, and event frequency, did not significantly differ between HC, AD APOE ε3/ε3, and AD APOE ε4/ε4 astrocytes, either under basal or mechanically stimulated conditions. These results indicate that Ca²⁺ signaling dynamics are preserved across the different astrocyte cultures, further supporting the functional competence of the differentiated cells.

After confirming two essential functional characteristics of mature astrocytes, we proceeded to investigate the astrocytic inflammatory response to further validate their functionality and evaluate the influence of the *APOE* genotype on reactivity to proinflammatory stimuli. The expression of a panel of proinflammatory and anti-inflammatory cytokine genes — *IL6, IL1B, TNFA, IL1A, IL2,* and *IL10* — was analyzed by RT-qPCR both in their basal state and after stimulation for 48 h with IL-1β + TNFα (Figure 3C, Figure 4, Supplementary Figure S2). Astrocytes from all genotypes demonstrated the ability to trigger a transcriptional inflammatory (*IL6, IL1B, TNFA, IL1A*) and anti-inflammatory (*IL10*) response upon stimulation (\**p*<0.05, \*\**p*<0.01, \*\*\**p*<0.001, \*\*\*\**p*<0.0001), with *IL2* consistently showing no response across all groups. Representative bright-field images illustrate the morphological transition from a flattened basal state to a more stellated appearance following inflammatory stimulation (Figure 3D). Additionally, to identify *APOE* genotype-specific variations in inflammatory gene expression, a wider array of genes including *NOS2*, *C3, CXCL3,* and *CHI3L1,* directly involved in innate immunity, was examined in the patient-derived and isogenic *APOE* cultures separately (Supplementary Figure S2, Figure 4). In patient-derived cultures, *C3, CXCL3,* and *CHI3L1* expression increased after the stimulus (\**p*<0.05, \*\**p*<0.01, \*\*\**p*<0.001, \*\*\*\**p*<0.0001), although no significant *APOE* genotype differences were observed for any gene under either condition (Supplementary Figure S2). However, in isogenic cultures, while all genes tested (except for *IL2* and *NOS2*) showed increased expression after IL-1β + TNFα treatment (\**p*<0.05, \*\**p*<0.01, \*\*\**p*<0.001, \*\*\*\**p*<0.0001), several genotype-specific differences emerged when compared to the APOE 3/3 basal condition (Figure 4). *CXCL3* expression was significantly higher under basal conditions in APOE 4/4 and APOE KO astrocytes compared to APOE 2/2 and APOE 3/3 (^#^*p*<0.05, ^##^*p*<0.01), suggesting an increased basal chemokine tone in the absence of functional APOE signaling. Furthermore, *IL6* mRNA levels were significantly elevated in APOE 4/4 astrocytes compared to APOE 3/3 under stimulated conditions (^#^*p*<0.05), while *IL10* exhibited increased basal expression in APOE KO relative to APOE 3/3 astrocytes (^#^*p*<0.05) (Figure 4). These findings reveal that although the overall inflammatory response capacity is preserved across genotypes, specific transcriptional differences become apparent in isogenic APOE 4/4 astrocytes under basal and stimulated conditions. Moreover, *CXCL3* expression may be modulated by APOE 4/4 through a loss-of-function mechanism as a similar upregulation was observed in KO astrocytes.

Given the prominent role of IL-6 as a crucial mediator of astrocytic neuroinflammation [75,76], its protein-level release was subsequently measured at 75 DIV using ELISA in both patient-derived and isogenic *APOE* astrocytes under both basal and stimulated conditions (IL-1β + TNFα) (Figure 3E). In AD patient-derived cultures, the IL-6 release significantly increased after inflammatory stimulation across all groups — HC, AD APOE 3/3 and AD APOE 4/4 (\*\*\*\**p*<0.0001). Interestingly, under basal conditions, AD APOE 4/4 astrocytes showed significantly higher IL-6 release compared to AD APOE 3/3 (^#^*p*<0.05), indicating a heightened basal inflammatory state specifically linked to the *APOE4* homozygous condition in the disease context. To assess whether this effect was due to the *APOE* genotype itself, IL-6 release was also examined in isogenic *APOE* cultures, where all genotypes showed a significant increase upon stimulation (\*\*\*\**p*<0.0001). Under basal conditions, APOE 4/4 astrocytes released significantly more IL-6 than all other genotypes (∼2.7-fold higher; ^#^*p*<0.05, ^##^*p*<0.01, ^####^*p*<0.0001), supporting a genotype-specific proinflammatory basal state independent of the disease background. Paradoxically, under stimulated conditions, APOE 4/4 astrocytes released significantly less IL-6 than APOE 3/3 cells (^#^*p*<0.05), suggesting that despite their elevated basal inflammatory tone, APOE 4/4 astrocytes exhibit an attenuated inducible IL-6 secretory response when faced with proinflammatory stimulation.

### APOE 4/4 astrocytes display a reduced basal cell size and increased branching upon inflammatory stimulation

Given that APOE 4/4 astrocytes display a dysregulated inflammatory profile, we next examined whether these alterations were reflected in morphological changes. As the preceding results pointed to *APOE* genotype as the primary driver of the observed astrocytic alterations, we focused subsequent experiments on isogenic *APOE* cultures to discern the specific contribution of each *APOE* allele. Due to inter-experimental variability, all quantitative data were normalized to the APOE 3/3 condition, considered to be the neutral allele [77], unless specified otherwise. Astrocyte morphology was evaluated under basal conditions and after inflammatory stimulation with IL-1β + TNFα across all *APOE* genotypes. ICC images of GFAP-stained astrocytes under both conditions are shown in Figure 5A. In basal conditions (DMEM/N2 plus 0.1% BSA in water), APOE 4/4 astrocytes displayed a significantly smaller area compared to all other genotypes (^#^*p*<0.05, ^##^*p*<0.01, ^###^*p*<0.001), indicating a reduced cell size specifically linked to *APOE4* homozygosity (Figure 5B). This reduction was partially reflected in perimeter, which was also significantly smaller in APOE 4/4 compared to APOE 3/3 (^##^*p*<0.01) and APOE 2/2 (^##^*p*<0.01), although no significant difference was found relative to KO astrocytes (Figure 5C). Further analysis, combining basal conditions from both inflammatory and Aβ stimulation experiments (Figure 6), confirmed a significantly smaller area (vs APOE 2/2 and KO; \*\**p*<0.01, \**p*<0.05) and reduced perimeter (vs APOE 3/3 and APOE 2/2; \**p*<0.05) in APOE 4/4 astrocytes (Supplementary Figure S3A and S3B). Additionally, a significantly higher solidity was noted in APOE 4/4 astrocytes compared to APOE 3/3 and APOE 2/2 (\**p*<0.05, \*\**p*<0.01) (Supplementary Figure S3C). Collectively, these findings point to a smaller and more compact basal morphology specifically associated with *APOE4* homozygosity.

**Figure 5.**
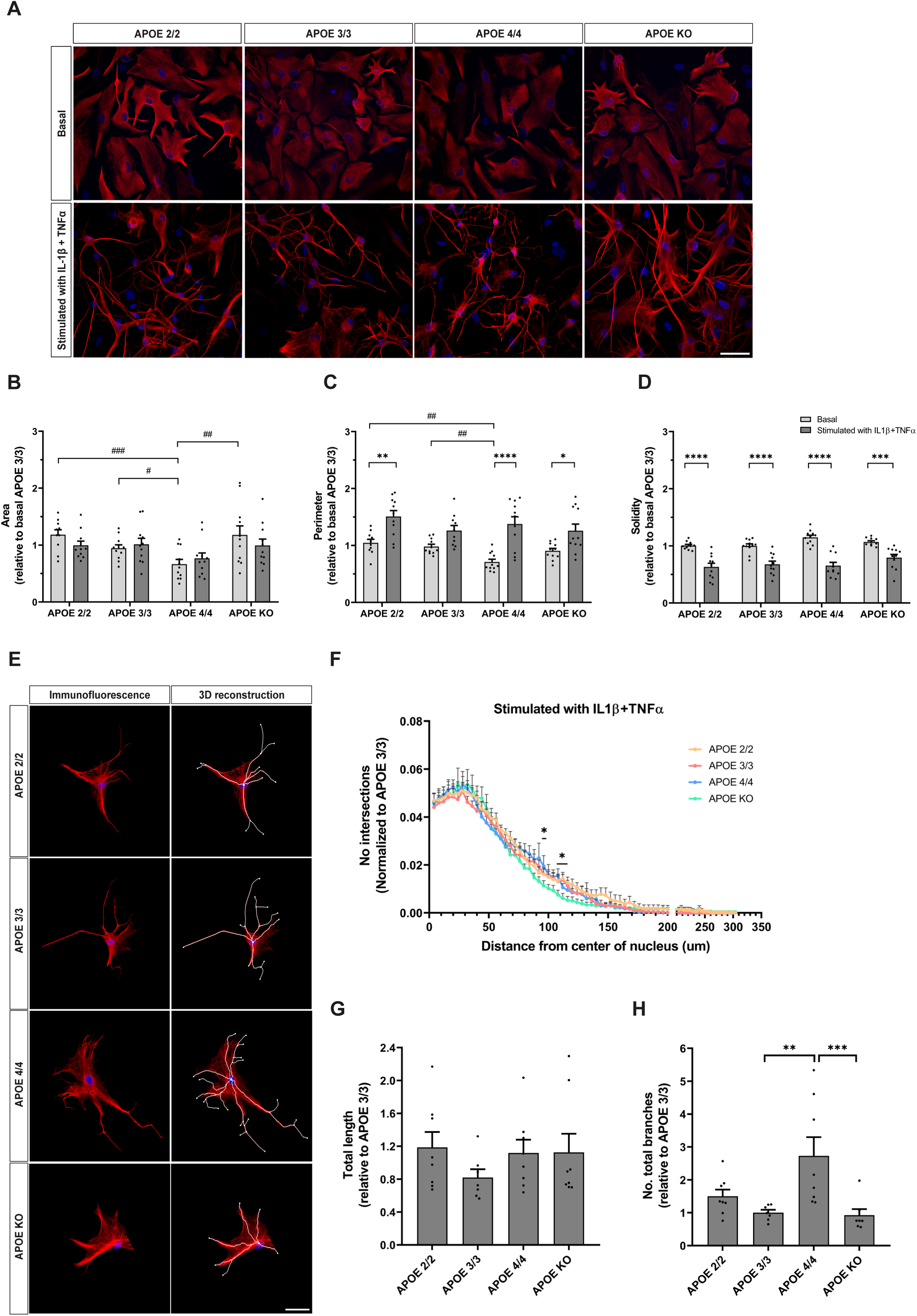
Proinflammatory stimulation unmasks a distinct morphological signature in APOE 4/4 astrocytes. **(A)** Representative immunofluorescence images of GFAP-stained astrocytes at 75-80 DIV under basal and proinflammatory conditions (IL-1β + TNFα) across *APOE* genotypes. Scale bar: 50 μm. **(B-D)** Morphometric profiling of area **(B)**, perimeter **(C)**, and solidity **(D)**. Under basal conditions, APOE 4/4 astrocytes exhibited significantly reduced area versus all genotypes, and smaller perimeter relative to APOE 2/2 and APOE 3/3. Stimulation did not alter area but significantly increased perimeter in APOE 2/2, APOE 4/4, and KO, with the strongest significant effect in APOE 4/4 cells. Solidity decreased significantly upon stimulation in all genotypes, with no between-genotype differences. Values were normalized to basal APOE 3/3 mean. Two-way ANOVA with Tukey’s test. Asterisks (*) indicate basal versus stimulated differences within genotype; hashtags (#) denote significant differences within condition. \**p*<0.05, ^#^*p*<0.05, \*\**p*<0.01, ^##^*p*<0.01, \*\*\**p*<0.001, ^###^*p*<0.001, \*\*\*\**p*<0.0001. Results are mean ± SEM of n = 10-12 independent cultures/ genotype. **(E)** Representative confocal images and 3D reconstructions of GFAP- positive astrocytes under stimulation, with digitally traced processes. Scale bar: 50 μm. **(F)** Sholl analysis under proinflammatory conditions, showing process intersections versus distance from the soma. APOE 4/4 and APOE 2/2 showed a minor increase in distal complexity versus APOE KO at 90–120 μm. Values were adjusted to APOE 3/3 mean. Two-way ANOVA with Tukey’s test. \*\**p*<0.01. Results are mean ± SEM of n = 8 independent cultures/genotype. **(G)** Total process length following proinflammatory stimulation, normalized to APOE 3/3 mean. No genotype differences were observed. One-way ANOVA with Tukey’s test. Results are mean ± SEM of n = 8 independent cultures/ genotype. **(H)** Total branch number following proinflammatory stimulation, normalized to APOE 3/3 value. APOE 4/4 exhibited significantly more branches than APOE 3/3 and APOE KO lines. One-way ANOVA with Tukey’s test. \*\**p*<0.01, \*\*\**p*<0.001. Results are mean ± SEM of n = 7-8 independent cultures/ genotype.

**Figure 6.**
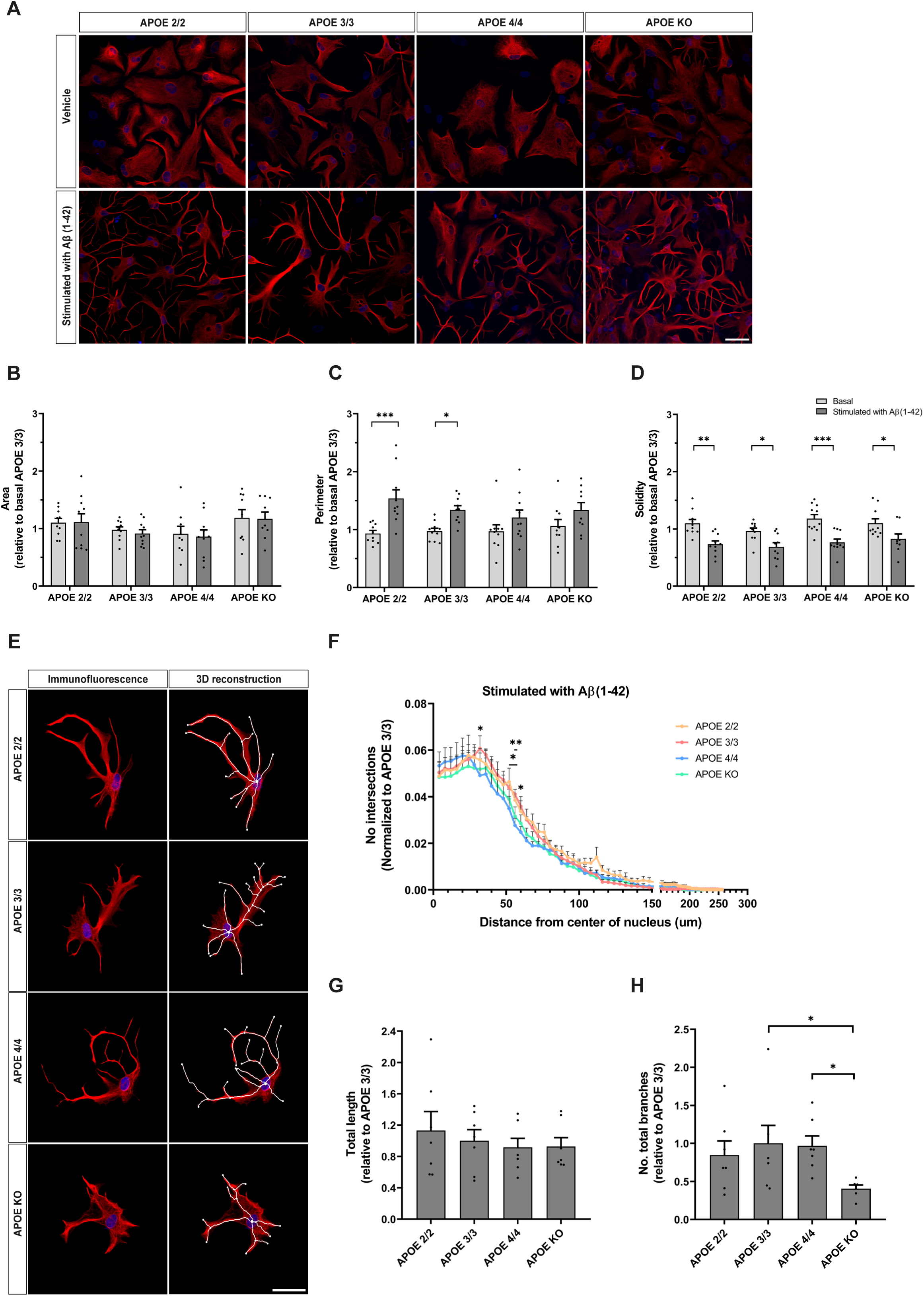
*APOE* genotype minimally influences astrocyte morphology under Aβ_1-42_ stimulation. **(A)** Illustrative GFAP immunofluorescence images of *APOE* isogenic astrocytes under vehicle and Aβ_1-42_ conditions at 75-80 DIV. Scale bar: 50 μm. **(B–D)** Morphometric profiling including area **(B)**, perimeter **(C)**, and solidity **(D)**. Area remained stable across genotypes and conditions. Aβ_1-42_ significantly increased perimeter in APOE 2/2, APOE 3/3, and APOE 4/4 astrocytes, with no inter-genotypic differences under vehicle conditions, and significantly decreased solidity across all genotypes. Values were normalized to vehicle APOE 3/3 mean. Two-way ANOVA with Tukey’s test. \**p*<0.05, \*\**p*<0.01, \*\*\**p*<0.001, \*\*\*\**p*<0.0001. Results are mean ± SEM of n = 10-12 independent cultures/ genotype. **(E)** Representative confocal images and 3D reconstructions of GFAP-stained astrocytes following Aβ_1-42_ exposure, with digitally traced processes. Scale bar: 50 μm. **(F)** Sholl analysis following Aβ_1-42_ treatment, showing process intersections versus distance from the soma. Branching complexity was significantly higher in APOE 2/2 and APOE 3/3 compared to APOE 4/4 cells at three isolated distances between 30-70 µm. Values were normalized to APOE 3/3 mean. Two-way ANOVA with Tukey’s test. \**p*<0.05, \*\**p*<0.01. Results are mean ± SEM of n = 7 independent cultures/ genotype. **(G)** Total process length upon Aβ_1-42_ treatment, normalized to APOE 3/3 mean. Cumulative process length was consistent across all genotypes. One-way ANOVA with Tukey’s test. Results are mean ± SEM of n = 7 independent cultures/ genotype. **(H)** Total branch number following Aβ_1-42_ treatment, relative to APOE 3/3 mean. APOE KO showed significantly fewer branches than APOE 3/3 and APOE 4/4 counterparts. One-way ANOVA with Tukey’s test \**p*<0.05. Results are mean ± SEM of n = 7 independent cultures/ genotype.

Upon inflammatory stimulation, the cell area remained consistent across all genotypes, with no differences observed between them (Figure 5B). However, the perimeter increased significantly in APOE 2/2 (\*\**p*<0.01), APOE 4/4 (\*\*\*\**p*<0.0001), and KO (\**p*<0.05) astrocytes (Figure 5C), while solidity decreased significantly across all genotypes (\*\*\**p*<0.001, \*\*\*\**p*<0.0001), with no differences detected between them (Figure 5D). These results suggest that despite their distinct basal morphology, APOE 4/4 astrocytes maintain a similar capacity for morphological remodeling in response to inflammatory stimuli, reflecting the typical transition towards a stellate reactive morphology (Figure 5A).

To thoroughly investigate whether the complexity of morphological changes differed among genotypes, Sholl analysis was conducted on reconstructed astrocytes under stimulated conditions. Figure 5E displays representative ICC images and 3D reconstructions. The overall Sholl profile revealed a broadly similar pattern of branching across genotypes (Figure 5F), with significant differences emerging only at discrete distances from the soma, as outlined in the pairwise comparisons in Supplementary Figure S4A-F. Specifically, APOE 2/2 and APOE 4/4 astrocytes exhibited significantly more branching intersections than APOE KO at two isolated points within the 90-120 μm range from the soma (\**p*<0.05) (Supplementary Figure S4C and S4F). Although total length did not significantly differ among genotypes, a non-significant trend towards shorter length was observed in APOE 3/3 astrocytes (Figure 5G). Conversely, the total branch number was significantly greater in APOE 4/4 astrocytes compared to APOE 3/3 (\*\**p*<0.01) and APOE KO (\*\*\**p*<0.001) cells (Figure 5H), despite the absence of differences in primary branches between genotypes (Supplementary Figure S3D). Together, these results suggest a genotype-specific morphological response to inflammatory stimulation in APOE 4/4 astrocytes, with an increased branching complexity without a corresponding increase in process length.

### Aβ_1-42_ stimulation induces a broadly comparable astrocyte morphological response across *APOE* genotypes

After determining that APOE 4/4 astrocytes exhibit a partially distinct morphological profile and branching pattern when stimulated with IL-1β + TNFα, we next asked whether these changes were specific to an inflammatory context or could also be triggered by Aβ_1-42_, a key pathological factor in Alzheimer’s disease [78]. To explore this, astrocyte morphology was evaluated in isogenic *APOE* cultures treated with Aβ_1-42_, using vehicle-treated cells (DMEM/N2 plus 5.8% DMSO in Tris Base) as a control (Figure 6). Representative ICC images of GFAP-stained astrocytes under both conditions are shown in Figure 6A.

Under vehicle conditions, no significant differences in area, perimeter, or solidity were observed between genotypes (Figure 6B-D). Upon Aβ_1-42_ stimulation, the cell area remained unchanged across all genotypes, while the perimeter significantly increased in APOE 2/2 (\*\*\**p*<0.001) and APOE 3/3 (\**p*<0.05) astrocytes, with a tendency towards an increase in the remaining genotypes (Figure 6B-C). This perimeter expansion was accompanied by a significant reduction in solidity across all genotypes (\**p*<0.05, \*\**p*<0.01, \*\*\**p*<0.001), with no differences detected among them (Figure 6D). These findings collectively indicate that Aβ_1-42_ stimulation prompts a morphological shift towards a reactive phenotype in all genotypes, with no major differences.

To further explore genotype-specific differences in process complexity and arborization, Sholl analysis was conducted on reconstructed astrocytes under Aβ_1-42_ stimulation, with representative ICC images and 3D reconstructions shown in Figure 6E. Although the overall Sholl profile was largely similar across genotypes (Figure 6F), pairwise comparisons revealed significant differences at specific points within the proximal 30-70 μm range from the soma (Figure 6F, Supplementary Figure S4G-L). APOE 2/2 astrocytes showed a significant difference at one point compared to APOE 4/4 (\**p*<0.05), with a trend towards increased branching throughout the entire 30-70 μm range (Supplementary Figure S4H). Similarly, APOE 3/3 astrocytes exhibited significantly higher branching intersections compared to APOE 4/4 at three points within this region (\**p*<0.05, \*\**p*<0.01), with a comparable trend (Supplementary Figure S4J). No differences were found between other genotype comparisons, suggesting a relative pattern of reduced proximal arborization specifically in APOE 4/4 cells upon Aβ_1-42_ stimulation. Regarding overall process metrics, total length was similar across genotypes (Figure 6G), while APOE KO astrocytes had a significantly lower total branch number compared to APOE 3/3 and APOE 4/4 cells (\**p*<0.05) (Figure 6H), with no differences in primary branches between genotypes (Supplementary Figure S3E). Together, these findings suggest a trend towards reduced proximal branching complexity in APOE 4/4 astrocytes upon Aβ_1-42_ stimulation, indicating a potentially distinct genotype-specific morphological response to amyloid pathology. Additionally, the absence of *APOE* partially modifies the astrocyte branching pattern in response to Aβ_1-42._

To investigate whether the differential response was influenced by the type of stimulus, Sholl profiles, total length, and total branch number were directly compared between IL-1β + TNFα and Aβ_1-42_ stimulation for each genotype (Supplementary Figure S5). The Sholl profiles of APOE 2/2 and APOE KO astrocytes were similar across both stimuli, indicating a similar level of morphological changes regardless of the stimulus nature (Supplementary Figure S5A, S5D). Conversely, APOE 3/3 astrocytes displayed significantly more branching intersections at three discrete points (\*\**p*<0.01, \*\*\*\**p*<0.0001) within the proximal 20-60 μm range under IL-1β + TNFα compared to Aβ_1-42_, with a tendency towards decreased complexity in more distal regions (70-140 μm) under Aβ_1-42_ (Supplementary Figure S5B). APOE 4/4 astrocytes showed a non-significant trend of increased branching complexity under IL-1β + TNFα compared to Aβ_1-42_ in the 60-110 μm range (Supplementary Figure S5C), along with a significantly higher total branch number under inflammatory conditions (\*\*\**p*<0.001) (Supplementary Figure S5F). Moreover, upon inflammatory stimulation, APOE 4/4 astrocytes showed significantly more branches than all other genotypes (^##^*p*<0.01; ^###^*p*<0.001). In contrast, the total length was consistent across genotypes and stimuli (Supplementary Figure S5E). These results collectively indicate that APOE 2/2 and APOE KO astrocytes react similarly to both stimuli, whereas APOE 3/3 and APOE 4/4 cells exhibit more pronounced morphological changes under IL-1β + TNFα stimulation, suggesting a stimulus-dependent and genotype-specific pattern of astrocytic morphological reactivity.

### *APOE4* homozygous astrocytes exhibit increased Aβ_1-42_ retention and a reduced basal calnexin-tagged ER compartment

Given the well-established role of astrocytes in Aβ clearance and the evidence that *APOE* genotype influences this capacity (de Leeuw et al., 2022; Lin 2018; Koutsodendris 2022; Xia 2024; Shostak 2026), we next explored whether *APOE4* homozygosity influences the capacity of isogenic astrocytes to internalize exogenously applied Aβ_1-42_, and its subsequent intracellular trafficking towards the endoplasmic reticulum (ER). Labeled Aβ_1-42_ was applied to isogenic *APOE* cultures, and its uptake was assessed by measuring the percentage of Aβ- positive astrocytes and the relative Aβ-tagged area per cell (Figure 7). The percentage of Aβ-positive astrocytes was significantly higher in APOE 4/4 compared to APOE KO cells (\**p*<0.05), with no statistically significant differences observed among the remaining genotypes (Figure 7A). However, the relative Aβ- tagged area per cell was similar across all genotypes (Figure 7B-C).

**Figure 7.**
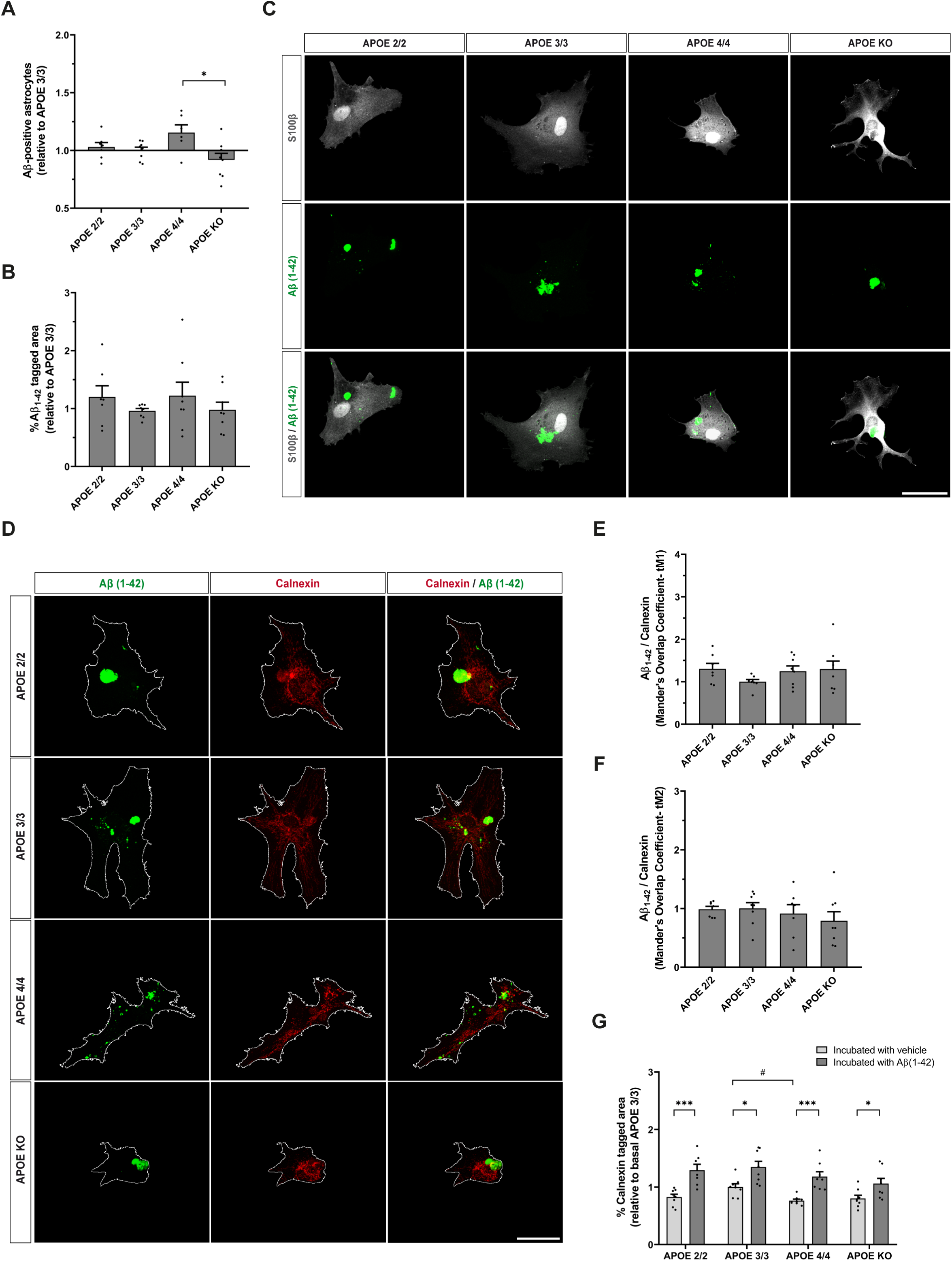
APOE 4/4 astrocytes show higher proportion of Aβ_1-42_-positive cells and reduced basal calnexin-tagged ER area. **(A)** Relative abundance of Aβ_1-42_ -positive astrocytes at 75-85 DIV across *APOE* isogenic lines, normalized to APOE 3/3. The proportion of Aβ_1-42_ -positive cells was significantly higher in APOE 4/4 astrocytes relative to the KO line, with no significant differences among the remaining genotypes. One-way ANOVA with Tukey’s test \**p*<0.05. Results are mean ± SEM of n = 7-8 independent cultures/ genotype. **(B)** Percentage of cellular area occupied by internalized Aβ_1-42_, normalized to APOE 3/3 mean. No genotype differences were observed. Kruskal-Wallis test with Dunn’s test. Results are mean ± SEM of n = 7-8 independent cultures/ genotype. **(C)** Representative fluorescence images showing Aβ_1-42_ distribution within S100β-labeled *APOE* isogenic astrocytes. Scale bar: 50 μm. **(D)** Representative confocal panels showing spatial distribution and potential colocalization of internalized Aβ_1-42_ with the ER marker calnexin across *APOE* genotypes. Scale bar: 50 μm. **(E-F)** Colocalization analysis using thresholded Mander’s overlap coefficients, representing the fraction of Aβ_1-42_ signal coincident with ER (tM1, **E**) and the fraction of ER area containing Aβ_1-42_ (tM2, **F**). No genotype differences were observed. One-way ANOVA with Tukey’s test or Brown-Forsythe and Welch ANOVA. Results are mean ± SEM of n = 7-8 independent cultures/ genotype. **(G)** Percentage of calnexin-positive area under basal and Aβ_1-42_ stress conditions, relative to basal APOE 3/3 mean. Aβ_1-42_ triggered a significant increase in calnexin-positive area across all genotypes, while APOE 4/4 showed significantly smaller calnexin-positive area than APOE 3/3 under basal conditions. Two-way ANOVA with Tukey’s test. Asterisks (*) indicate vehicle versus Aβ_1-42_ differences within genotype; hashtags (#) denote genotype differences between within condition. \**p*<0.05, ^#^*p*<0.05, \*\*\**p*<0.001. Results are mean ± SEM of n = 7-8 independent cultures/ genotype.

We then investigated the intracellular localization of Aβ_1-42_. Considering that APOE 4/4 is associated with ER stress and altered protein processing in astrocytes [50], we examined whether internalized Aβ_1-42_ was localized to the ER by quantifying its colocalization with calnexin, a known ER marker, using Mander’s overlap coefficients (Figure 7D-F). The colocalization was consistent across genotypes for both coefficients, suggesting that the localization of Aβ_1-42_ to the ER is not influenced by *APOE* alleles. However, analysis of the ER compartment itself revealed a genotype-dependent variation (Figure 7G). The percentage of calnexin-tagged area per cell increased significantly upon Aβ_1-42_ stimulation across all genotypes (\**p*<0.05; \*\*\**p*<0.001), indicating a structural ER response to amyloid exposure. Notably, APOE 4/4 astrocytes exhibited a significantly smaller basal calnexin-tagged area compared to APOE 3/3 (\**p*<0.05), suggesting a proportionally reduced basal ER compartment specifically linked to the homozygous ε4 state.

Overall, these results point towards a greater proportion of APOE 4/4 astrocytes retaining detectable Aβ_1-42_, although the extent per cell and its ER localization remain consistent across genotypes. Furthermore, the diminished basal calnexin- labeled ER compartment in APOE 4/4 astrocytes may contribute to a compromised protein processing capacity associated with the ε4 allele.

## Discussion

Here we have used isogenic and non-isogenic hiPSCs derived from AD patients to investigate the impact of *APOE* polymorphism on astrocyte maturation and function and its involvement in AD pathology. Using fully defined cell culture media, neural inducers and growth factors, we have generated hiPSC-derived astrocytes that recapitulate main astrocyte molecular, cellular, and functional features. These include molecular marker expression, cell morphology, intracellular Ca²⁺ responses, glutamate uptake capacity, cytokine and chemokine expression in response to inflammatory stimulus, and Aβ_1-42_ uptake capacity. We report that astrocytes carrying the ε4 allele in homozygosis (APOE 4/4 astrocytes) show increased basal IL-6 release and *CXCL3* expression, reduced glutamate uptake, and altered Aβ handling alongside reduced GFAP and S100β protein expression and a smaller, more compact basal morphology.

### APOE 4/4 impairs astrocyte maturation while promoting a basal proinflammatory and neurotoxic state

Here we show that both the percentage of GFAP and S100β-expressing astrocytes as well as cell body area were significantly reduced in APOE 4/4 cultures, indicating that the ε4 allele in homozygosis alters the acquisition of mature astrocytic features. Notably, this reduction was restricted to the protein level: *GFAP* and *S100B* transcripts increased during differentiation with no differences between genotypes, revealing a dissociation between mRNA and protein synthesis for these two independent markers. Therefore, this alteration appears downstream of transcription and is consistent with evidence that *APOE4* compromises proteostasis in glial cells [48,79]

Independently of these maturation phenotypes, APOE 4/4 promotes an innate proinflammatory state in the absence of an exogenous inflammatory stimulus, suggesting a chronic, basal priming of the astrocytic proinflammatory response. This is supported by our results showing that APOE 4/4 increases IL-6 release and *CXCL3* expression compared to other APOE genotypes, two key components of the innate inflammatory activation [80]. In support of the idea that our culture system recapitulates key features of APOE 4/4-AD, higher circulating levels of IL-6 were observed in APOE 4/4 human carriers [81–83].

Our results suggest that APOE 4/4-mediated processes are independent of cell size and of GFAP and S100β expression and indicate that these cellular features are not critical for the induction of the basal proinflammatory state by APOE 4/4. In addition, APOE 4/4 reduces astrocyte glutamate uptake from the medium. If this effect were to correlate with the *in vivo* situation, the milieu surrounding neurons and glial cells could contain elevated glutamate concentrations, potentially triggering neurotoxicity [84]. Altogether, our results suggest that APOE 4/4 compromises brain homeostasis by promoting an inflammatory and neurotoxic environment. Although these features were partially observed in non-isogenic astrocytes derived from APOE 3/3 and APOE 4/4 AD patients compared to healthy controls, the use of isogenic hiPSCs strengthened the case for APOE 4/4-specific phenotypes.

Under stimulation conditions, IL-1β + TNFα strongly triggers IL-6 release from APOE 2/2, APOE 3/3, APOE 4/4 and APOE KO astrocytes. Under these conditions, APOE 4/4 astrocytes have significantly more branches, although they release significantly less IL-6 than APOE 3/3 astrocytes, again suggesting that the morphological and secretory components of the astrocytic response to IL-1β + TNFα may partially function as independent processes. By contrast, *IL6* mRNA expression in APOE 4/4 astrocytes was higher than in APOE 3/3 astrocytes, suggesting a mismatch between transcription and secretion of IL-6 upon IL-1β + TNFα treatment.

Previous studies reported a basal inflammatory phenotype of APOE 4/4 astrocytes, in part characterized by increased IL-6 release and higher *IL6* mRNA expression [43,45,49,85]. Our new findings show that this increase is functionally dissociated from changes in cell size, calnexin-tagged ER area as well as from GFAP and S100β expression, as mentioned above. However, they suggest a potential correlation between Aβ accumulation and IL-6 synthesis and release in APOE 4/4 astrocytes under basal conditions. This idea is supported by the increased proportion of Aβ-positive APOE 4/4 astrocytes compared to other APOE genotypes, a difference that reached statistical significance versus APOE KO astrocytes. This view is in line with previous studies reporting that APOE 4/4 promotes Aβ aggregation and impairs Aβ clearance, inducing inflammation [21,29,83,86].

Another relevant observation to protein handling is the reduced basal calnexin- tagged ER compartment in APOE 4/4 astrocytes, which cannot be explained by the smaller size of APOE 4/4 cells because this measure was expressed as a percentage of cell area. Aβ_1-42_ exposure expanded the calnexin-positive compartment similarly in all genotypes, indicating than APOE 4/4 astrocytes retain the capacity to generate an ER response to a toxic challenge but do so from a significantly reduced baseline. A smaller resting ER is consistent with reports that APOE4’s domain-interaction properties trigger an unfolded protein response and impair astrocyte function in a mouse model [50], and offers a plausible structural correlate — though not proof — of the post-transcriptional and secretory abnormalities described above. Considered together, the mRNA/protein dissociation for GFAP and S100β, the mismatch between *IL6* transcription and IL-6 secretion, and the reduced ER compartment converge on the possibility that APOE 4/4 compromises astrocytic protein processing at several levels rather than affecting any single pathway in isolation.

The reduced glutamate uptake by APOE 4/4 astrocytes compared to APOE 3/3 is an additional key process that could contribute to dysregulated cellular homeostasis. Reduced uptake causes extracellular glutamate to accumulate, potentially increasing excitotoxic stress [84]. Thus, the basal inflammatory state observed in APOE 4/4 astrocytes could affect the expression or activity of glutamate transporters impairing glutamate uptake. In turn, high extracellular glutamate could influence neuroinflammation by stimulating intracellular signaling pathways leading to increased IL-6 production. Therefore, the astrocyte basal inflammatory state could be associated with impaired glutamate uptake and clearance contributing to a neurotoxic environment [87].

We show that APOE 4/4 astrocytes display a more compact basal morphology, together with an exaggerated branching response to IL-1β + TNFα that was not observed with Aβ_1-42_, indicating a stimulus-specific divergence between inflammatory and amyloid challenge. This suggests that APOE 4/4 does not make astrocytes more or less morphologically reactive, but instead determines which stimuli elicit a morphological response. Supporting this, the direct comparison of both stimuli within the same genotype confirmed that APOE 4/4 astrocytes branch more under inflammatory than under amyloid challenge, a divergence not observed in the other genotypes. These findings are consistent with growing evidence that astrocyte reactivity is not a single program but a heterogeneous response shaped by the nature of the eliciting stimulus, both at the transcriptional [88] and morphological levels (Escartin et al., 2021), and with calls to move away from binary reactivity categories toward multi-parameter assessment that includes morphology alongside molecular and functional readouts.

Our results are partially in line with Preman et al. (2021), who reported that astrocyte morphological changes around amyloid plaques were largely independent of *APOE* genotype when comparing APOE 3/3 and APOE 4/4. Using a complete allelic series including APOE 2/2 and APOE KO, we extend this by showing that branching distribution (number of intersections), rather than cell shape (area, perimeter, and solidity), is where genotype-dependent differences in response to Aβ_1-42_ may emerge.

### APOE 4/4 largely acts through gain-of-function mechanisms in human astrocytes

A number of studies have reported that humanized *APOE4* mice exhibit multiple gain-of-function phenotypes, such as synaptic dysfunction, altered lipid metabolism, mitochondrial impairment, altered glial activation, and enhanced tau toxicity. Importantly, some phenotypes have been validated in post-mortem human samples and may arise independently of, or in association with, amyloid deposition. Therefore, lowering APOE 4/4 has emerged as a potential treatment for APOE ε4/ε4 carriers [83,89–93]. Our results also support a largely gain-of-function mechanism of *APOE4* in the observed phenotypes. Most of the phenotypes we describe here — reduced GFAP marker expression, smaller and more compact basal area, reduced glutamate uptake, elevated basal IL-6 release, increased branching upon cytokine stimulation, and increased number of Aβ-positive astrocytes — were not reproduced in APOE KO astrocytes. If these phenotypes simply reflected loss of a protective APOE function, KO astrocytes should show them at least as strongly as APOE 4/4 cells. However, this was not the case, indicating that APOE 4/4 protein itself must be present to drive a gain-of-toxic-function mechanism perturbing astrocyte homeostasis.

However, our findings show that the regulation by APOE 4/4 of molecular components of baseline innate immunity appears to involve both gain-of-function (IL-6 release and expression) and loss-of-function (*CXCL3* expression) mechanisms. This latter finding suggests that eliminating *APOE* gene function may favor *CXCL3* production by astrocytes. APOE KO astrocytes also showed significantly higher basal anti-inflammatory *IL10* expression, indicating that loss of *APOE* dysregulates basal cytokine and chemokine expression more broadly than a single proinflammatory shift. Therefore, additional research would be necessary to understand the impact of eliminating *APOE* on the activity of the innate immune system.

Importantly, converting the parental line from *APOE* ε4/ε4 to *APOE* ε2/ε2 or *APOE* ε3/ε3 rescues the astrocyte phenotypes caused by APOE 4/4, suggesting a potential alternative therapeutic approach for APOE 4/4-associated AD.

**In conclusion,** using a collection of both non-isogenic and isogenic hiPSCs, we have identified key features of astrocyte dysfunction in AD that may emerge during relatively early stages of astrocyte development. In this context, APOE 4/4 may specifically disrupt basal cellular homeostasis by creating a proinflammatory and toxic milieu with potential consequences for brain homeostasis during AD progression.

### Limitations section

A limitation of our study is the absence of a direct *in vivo* translation of the results found in cultured human astrocytes. However, the use of isogenic astrocytes strengthened the conclusion that APOE 4/4 is a major toxic variant that could be involved in AD etiopathology since an early preclinical stage.

## Data Availability

The data that support the findings of this study are available on reasonable request from the corresponding author.

## Acknowledgements

We are grateful to Dr. Oliver Brüstle (University Hospital Bonn, Germany) and Dr. Benjamin Schmid (Bioneer A/S, Denmark) for sharing with us the parental *APOE* ε4/ε4 line and the derived isogenic cell lines (*APOE* ε2/ε2, *APOE* ε3/ε3, and *APOE* KO) in the frame of the ADAPTED project. We also thank Mª José Román for her technical support, Marta González Martín and Mª José Román (Instituto Cajal, CSIC, Spain) for their help with the image analysis, and Dr. Alexander Rodero (Instituto Cajal, CSIC, Spain) for his participation in pilot studies. We also thank the “Unidad de Imagen Científica y Microscopía” (Instituto Cajal, CSIC, Spain) and “Unidad Master” (Centro de Investigaciones Interdisciplinares de Alcalá, Ci2A and CNC, CSIC, Spain) for helping us with the image analysis. We acknowledge support from PTI^+^ NEUROAGING, CSIC, Spain.

## Authors contributions

R.V.: Designed and performed cell culture experiments, cellular and molecular assays, data analysis and interpretation, manuscript writing and revision. E.D.- G.: Designed and performed cell culture experiments, cellular and molecular assays, data analysis and interpretation. E.A.-G.: Designed and performed cell culture experiments, cellular and molecular assays, data analysis and interpretation. D.S.G.: Performed immunocytochemistry, morphological analysis and data interpretation. F.J.F.A.: Performed cell culture experiments, cellular and molecular assays. I.S.: Performed Ca²⁺ imaging experiments, data analysis and interpretation. E.P.M.-J: Performed cell culture experiments, cellular and molecular assays. R.M.: Manuscript revision, financial and administrative support. M.N.: Experimental design of Ca²⁺ imaging experiments, data analysis and interpretation, manuscript revision, financial and administrative support. C.V.: Conceptualization, experimental design, cell culture supervision, data analysis and interpretation, manuscript writing, revision and editing, financial and administrative support, final approval of manuscript.

## Ethics approval and consent to participate

The study was approved by the Commission of Guarantees for the Donation and Utilization of Human Cells and Tissues, Instituto de Salud Carlos III (ISCIII), Spain (approval numbers: 3202691, 3973231, 4093321, 4083311, and 5224261) and by the CSIC Ethics Committee, Spain (approval number 256/2023). Informed consent was obtained from all the patients.

