## Supplemental Table 1, Supplemental Figure Legends, Supplemental Figures for "APOE 4/4 promotes dysfunctional and inflammatory phenotypes concomitant with impaired maturation of hiPSC-derived astrocytes"

| Table S1. Primers (5’-3’) used for the gene-expression analysis by RT-qPCR | | | |
|---|---|---|---|
| *APOE* | Forward | CCAATCACAGGCAGGAAGATG | 55 bp |
| | Reverse | GGAATGTGACCAGCAACGCA | |
| *AQP4* | Forward | CATTGGAGCAGGAATCCTCTATCTGG | 176 bp |
| | Reverse | TGACATCAGTCCGTTTGGAATCACA | |
| *C3* | Forward | GAACTGCCTTTGTCATCTTC | 183 bp |
| | Reverse | CAGACACGTACAAAGACTTC | |
| *CHI3L1* | Forward | TGTACCCACATCATCTACAG | 137 bp |
| | Reverse | ACAGACAAGAGAGTCTTCAG | |
| *CXCL3* | Forward | CCTCAAGAACATCCAAAGTG | 154 bp |
| | Reverse | CCCCTTGTTCAGTATCTTTTC | |
| *GAPDH* | Forward | AACCATGAGAAGTATGACAACAGCC | 210 bp |
| | Reverse | TGAGTCCTTCCACGATACCAAAGT | |
| *GFAP* | Forward | GAGGCAGAAGCTCCAGGATGAAAC | 137 bp |
| | Reverse | TCTCCTCCTCCAGCGACTCAATCT | |
| *SLC1A2 (GLT1)* | Forward | TCATCCTGGGAGCAGTGTGTGG | 238 bp |
| | Reverse | ACTGCAGCAATGATGGTCGTGG | |
| *IL1A* | Forward | AGAGGAAGAAATCATCAAGC | 122 bp |
| | Reverse | TTATACTTTGATTGAGGGCG | |
| *IL1B* | Forward | CCTGAGCTCGCCAGTGAAATGAT | 163 bp |
| | Reverse | TGCTGTAGTGGTGGTCGGAGATTC | |
| *IL2* | Forward | CACTAAGTCTTGCACTTGTC | 158 bp |
| | Reverse | CTTAAATGTGAGCATCCTGG | |
| *IL6* | Forward | TGTGTGAAAGCAGCAAAGAGGCA | 125 bp |
| | Reverse | ACCAGTGATGATTTTCACCAGGCA | |
| *IL10* | Forward | GCGCTGTCATCGATTTCTTCCC | 248 bp |
| | Reverse | TCAGCTATCCCAGAGCCCCAGA | |
| *NOS2* | Forward | TCTTCGAAATCCCACCTGACCTTG | 178 bp |
| | Reverse | TCTGTGCCCATGTACCAGCCATT | |
| *S100B* | Forward | AAGCACAAGCTGAAGAAATCCGAAC | 181 bp |
| | Reverse | AGGCAGTAGTAACCATGGCAACAAAG | |
| *TNFA* | Forward | GAACCCCGAGTGACAAGCCTGTAG | 123 bp |
| | Reverse | GGTTATCTCTCAGCTCCACGCCAT | |

### SUPPLEMENTARY FIGURE LEGENDS

**Supplementary Figure S1. Temporal maturation trajectories of astrocytic markers in non-isogenic and isogenic hiPSC-derived astrocytes (A)** Relative mRNA expression of astrocytic markers *GFAP*, *AQP4*, *S100B* and *APOE* in neural progenitors (8-16 DIV) and astrocytes from HC, AD APOE 3/3 and AD APOE 4/4 patients at 60 and 90 DIV. All three groups exhibited a progressive, time-dependent increase in marker expression during differentiation-maturation, reaching their highest expression levels at 90 DIV. Statistical analysis was performed on ΔCt values using one-way ANOVA with Tukey's test; data are presented as $\log_2$ fold change relative to progenitors. *$p$<0.05, **$p$<0.01, ***$p$<0.001, ****$p$<0.0001. Results are mean ± SEM from n = 3 independent cultures in technical triplicates. **(B)** Relative mRNA expression of astrocytic markers *AQP4*, *S100B*, *APOE* and *SLC1A2*/*GLT1* in astrocytes derived from isogenic *APOE* hiPSC lines (ε2/ε2, ε3/ε3, ε4/ε4 and KO) at 60–75 DIV relative to progenitors (8-16 DIV). Significant upregulation was observed for most markers at 75 DIV across all isogenic lines relative to the progenitor stage. Statistical analysis was performed on ΔCt values using an unpaired t-test; data are shown as $\log_2$ fold change. *$p$<0.05, ***$p$<0.001, ****$p$<0.0001. Results are mean ± SEM from n = 4 independent cultures in technical triplicates. Together, these within-genotype maturation trajectories validate consistent astrocytic differentiation across all lines, complementing the between-genotype comparisons presented in Figures 1D and 2C. AD, Alzheimer's disease; HC, healthy controls.

**Supplementary Figure S2. Cytokine profiling shows conserved inflammatory response across HC and AD genotypes in non-isogenic human astrocytes.** mRNA expression of an extended panel of cytokines and inflammatory markers was analyzed in human astrocytes (75-90 DIV) from HC and AD patients following stimulation with IL-1β + TNFα, complementing the core cytokine panel shown in Figure 3C. No significant differences were observed between genotypes under either basal or stimulated conditions for any of the genes analyzed. Proinflammatory stimulation significantly upregulated *IL6*, *TNFA, IL10, C3, CXCL3*, and *CHI3L1* across all groups. For *IL1A, IL1B* and *NOS2*, the stimulation-induced increase reached statistical significance only in a subset of genotypes, while remaining directionally consistent across groups. No significant induction was detected for *IL2* in any group. Statistical significance was assessed on ΔCt values by two-way ANOVA with Tukey's test; data are expressed as $log_2$ fold change relative to the HC basal group. **p*<0.05, ***p*<0.01, ****p*<0.001, *****p*<0.0001. Results are mean ± SEM of n = 4 independent cultures/ genotype. AD, Alzheimer's disease; HC, healthy controls.

**Supplementary Figure S3. Pooled baseline data point to a smaller, more compact basal morphology in APOE 4/4 astrocytes. (A-C)** Cell area **(A)**, perimeter **(B)**, and solidity **(C)** were assessed using pooled baseline and vehicle data from the experimental sets shown in Figures 5B-D and 6B-D, normalized to APOE 3/3 mean value. APOE 4/4 astrocytes exhibited reduced cell area relative to APOE 2/2 and APOE KO, reduced perimeter relative to APOE 2/2 and APOE 3/3, and increased solidity compared to both APOE 2/2 and APOE 3/3 lines. One-way ANOVA with Tukey's test **p*<0.05, ***p*<0.01. Results are mean ± SEM of n

= 20-22 independent cultures/ genotype. **(D-E)** Number of primary branches under IL-1β + TNFα **(D)** or A$β_{1-42}$ **(E)** treatment, relative to APOE 3/3 mean. No significant differences were detected among *APOE* isogenic lines following either stimulation. Kruskal–Wallis test with Dunn's test. Results are mean ± SEM of n = 7-8 independent cultures/ genotype.

**Supplementary Figure S4. Pairwise Sholl comparisons reveal discrete *APOE* genotype-specific branching differences upon inflammatory and A$β_{1-42}$ stimulation.** Sholl analysis of human isogenic *APOE* astrocytes stimulated with IL-1β + TNFα **(A-F)** or exposed to A$β_{1-42}$ **(G-L)** at 75-80 DIV, evaluating pairwise differences in process branching complexity between genotypes. This analysis complements the aggregate Sholl profiles shown in Figures 5F and 6F. Data were normalized to the APOE 3/3 mean. Under IL-1β + TNFα stimulation, APOE 2/2 and APOE 4/4 cells showed minor, isolated increases in branching complexity relative to the APOE KO group **(C, F)**. Following A$β_{1-42}$ exposure, APOE 4/4 astrocytes displayed a localized reduction in branching density compared to both APOE 2/2 and APOE 3/3 lines at discrete distances **(H, J)**. Two-way ANOVA with Tukey's test *$p<0.05$, **$p<0.01$. Results are mean ± SEM of n = 7-10 independent cultures/ genotype.

**Supplementary Figure S5. Pairwise Sholl comparisons between inflammatory and A$β_{1-42}$ stimulation reveal discrete genotype-specific branching differences in astrocytes. (A-D)** Comparative Sholl analysis of astrocytic process complexity in *APOE* isogenic astrocytes after IL-1β + TNFα or

A$\beta_{1-42}$ stimulation, relative to APOE 3/3 mean value. A divergence in branching complexity between stimuli was detected specifically in APOE 3/3 astrocytes, restricted to two consecutive distances within the 30-50 μm range, with no significant differences observed in the other genotypes.Two-way ANOVA with Tukey's test $^{**}p<0.01$, $^{****}p<0.0001$. Results are mean ± SEM of n = 7-10 independent cultures/ genotype and stimulus. **(E)** Total process length of human astrocytes with different *APOE* genotypes following IL-1β + TNFα or A$\beta_{1-42}$ treatment, relative to APOE 3/3 mean. No significant differences in total process coverage were detected between the stimuli within any genotype. Two-way ANOVA with Tukey's test. Results are mean ± SEM of n = 7-10 independent cultures/ genotype and stimulus. **(F)** Total branch number across *APOE* genotypes and stimuli, relative to APOE 3/3 mean. Branch number increased significantly in APOE 4/4 astrocytes under IL-1β + TNFα compared to both A$\beta_{1-42}$ treatment in the same genotype and all other genotypes under the same inflammatory condition. Two-way ANOVA with Tukey's test. Asterisks (*) indicate differences between stimuli within genotype; hashtags (#) denote genotype differences within the stimulus condition. $^{\#\#}p<0.01$, $^{***}p<0.001$, $^{\#\#\#}p<0.001$. Results are mean ± SEM of n = 7-10 independent cultures/ genotype and stimulus.

Supplementary Figure S1

### Non-isogenic human astrocytes

**A**

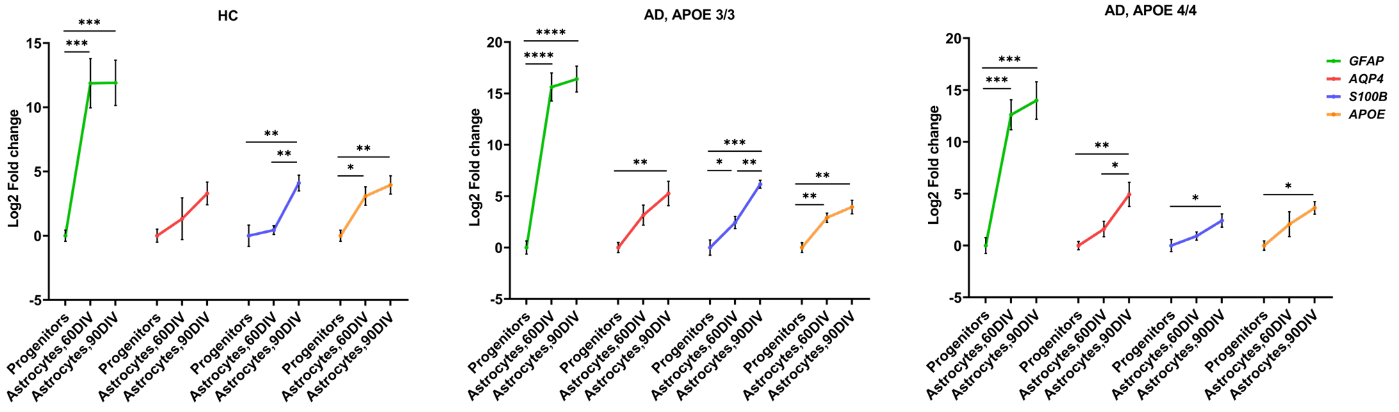


### Isogenic human astrocytes

**B**

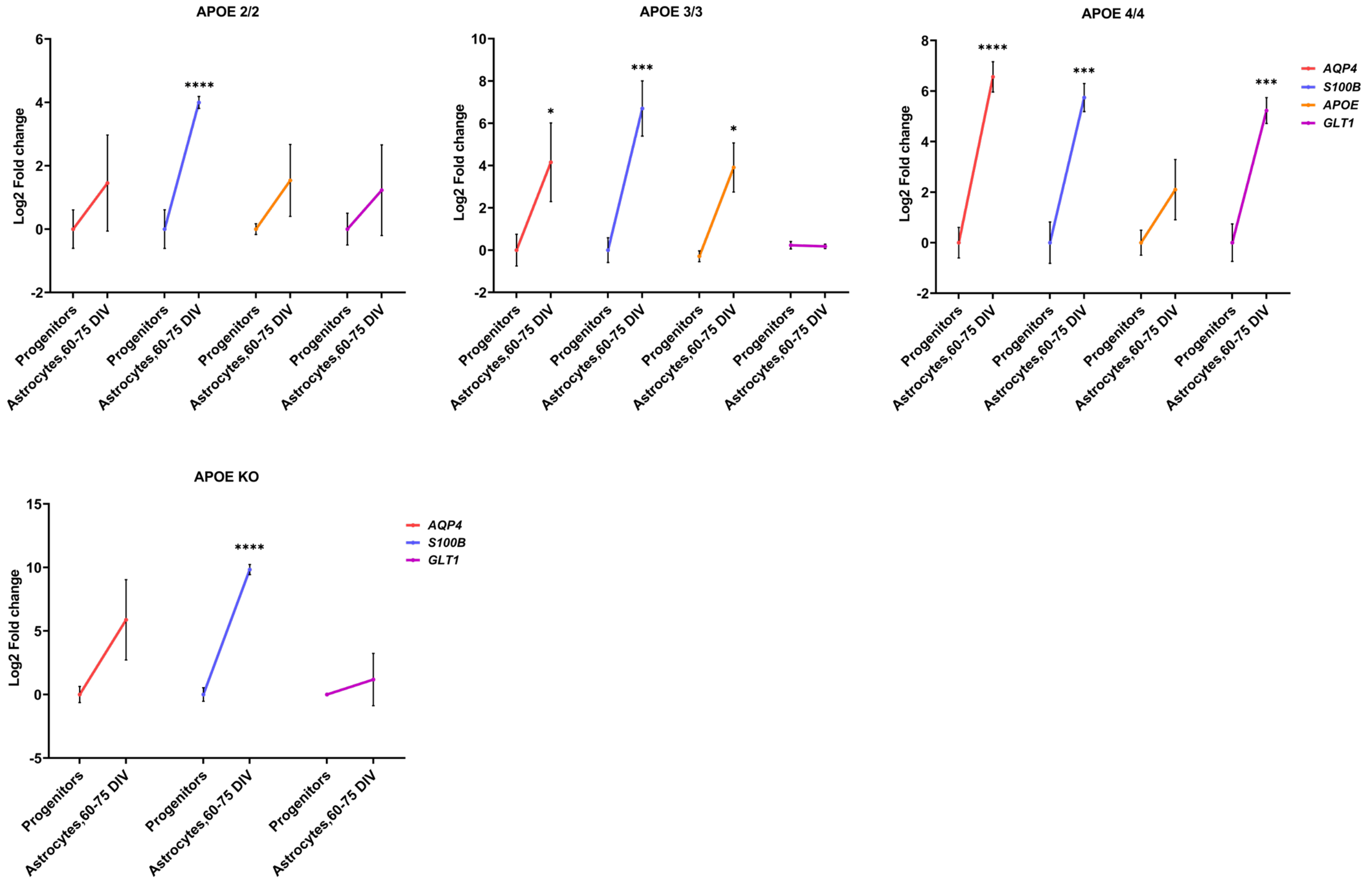

### Non-isogenic human astrocytes

**A**

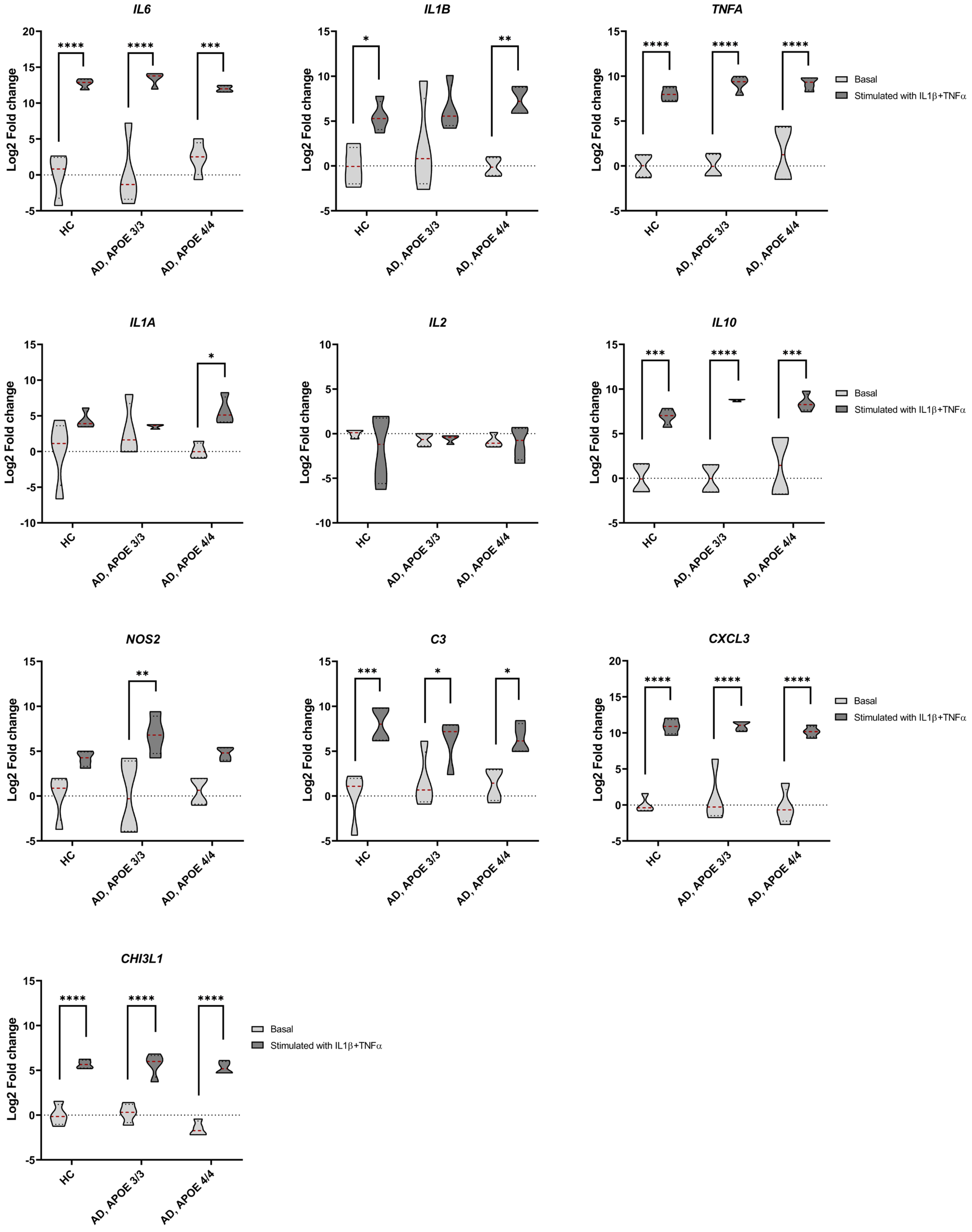

### Basal + vehicle conditions

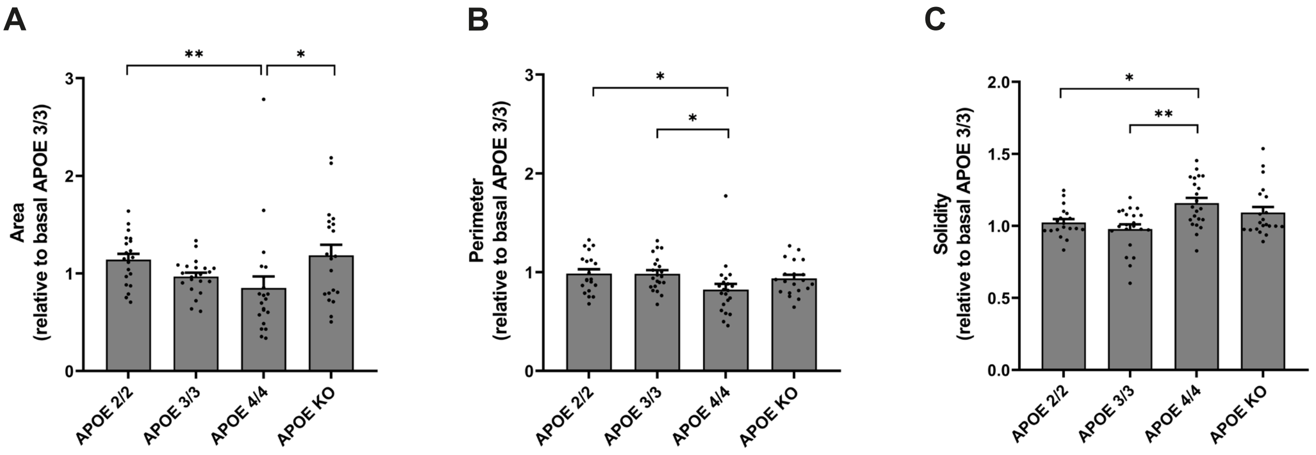


### Stimulated conditions: IL-1β + TNFα

### Stimulated conditions: Aβ (1-42)

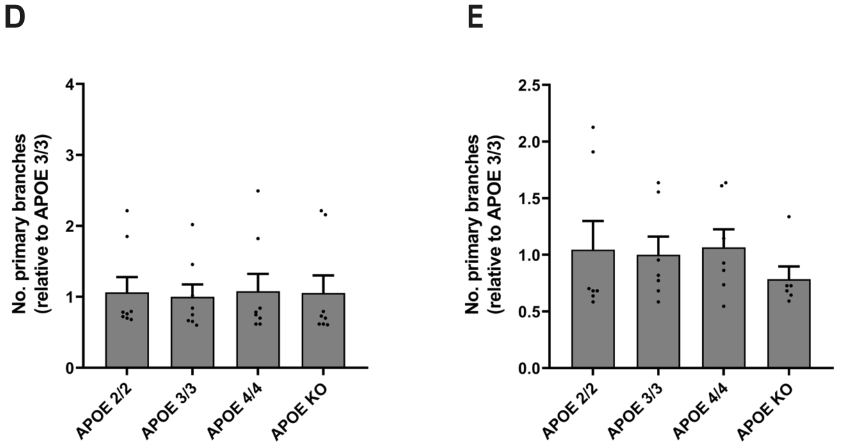

### Stimulated conditions: IL-1β + TNFα

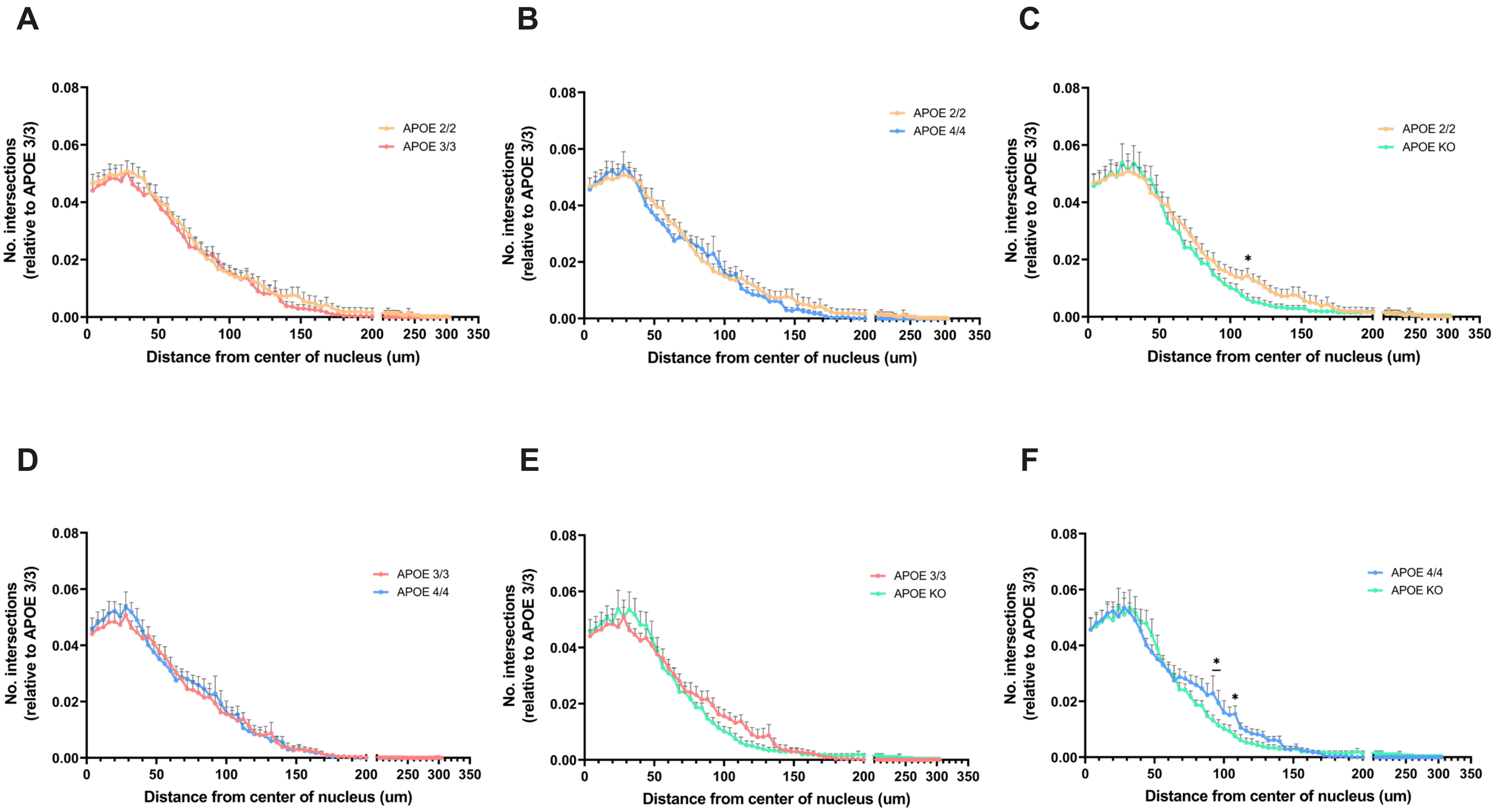


### Stimulated conditions: Aβ (1-42)

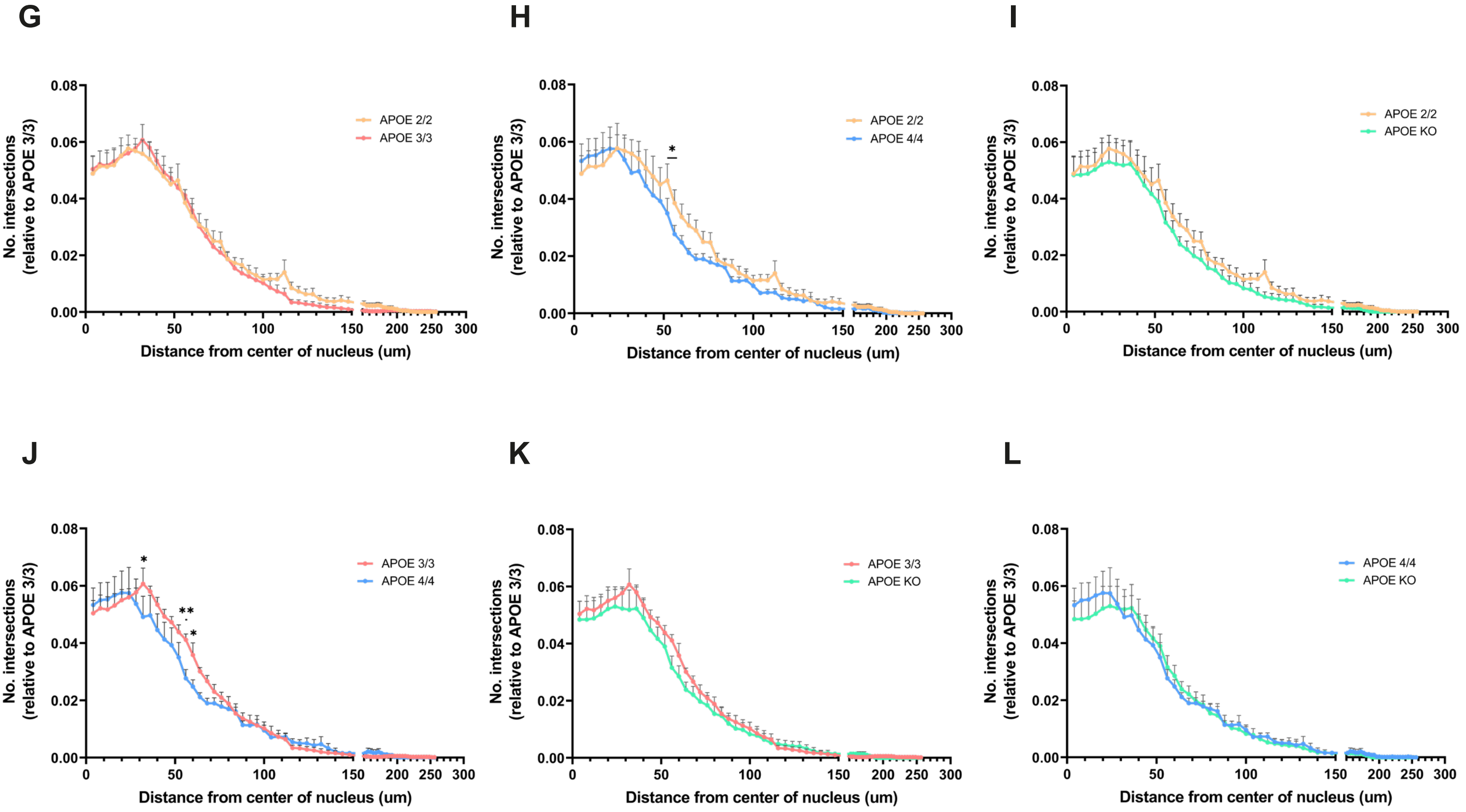

Supplementary Figure S5

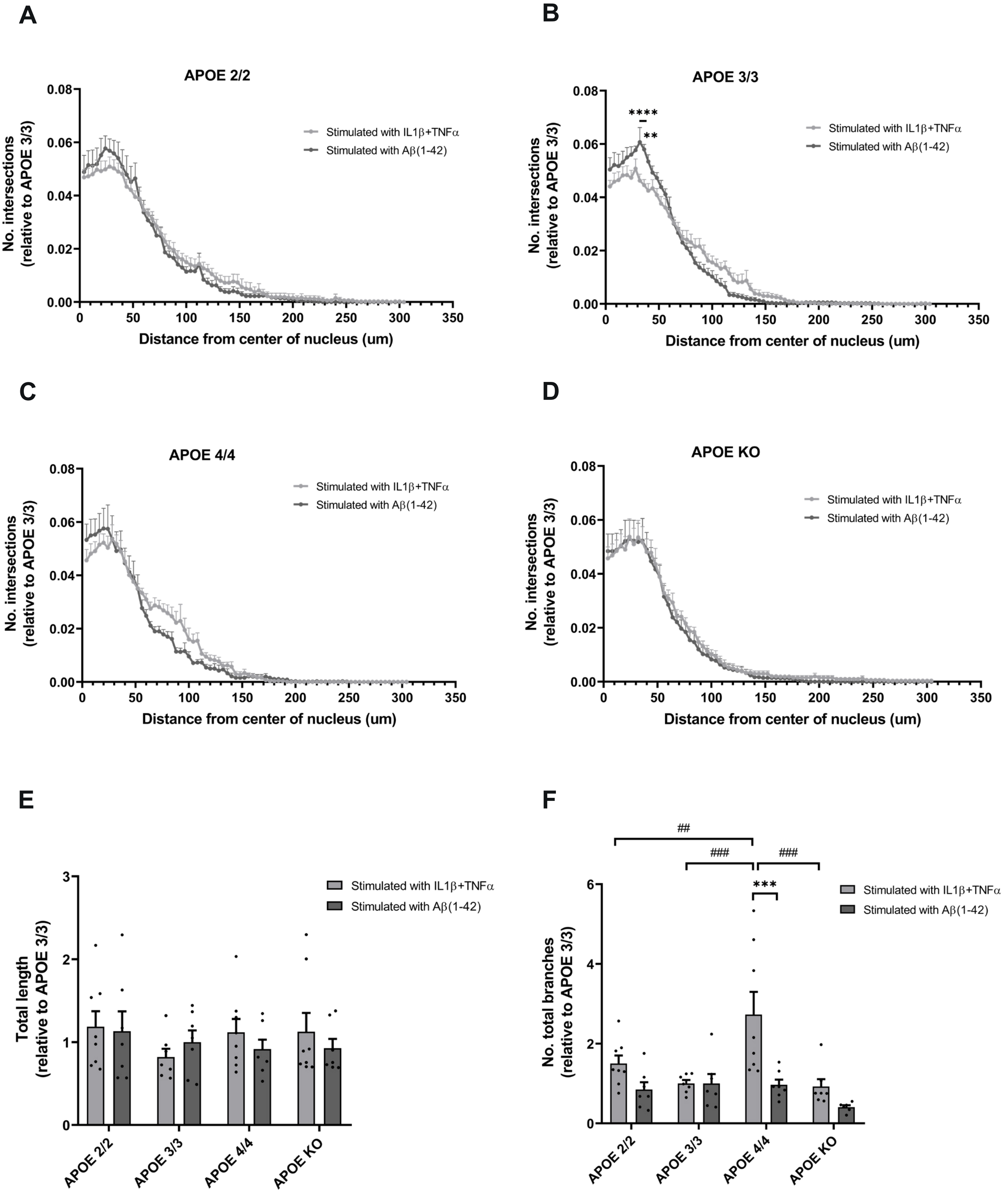
